# A distributed genomic architecture of repeated thermal adaptation in a low-dispersal marine invader

**DOI:** 10.64898/2026.09.28.755179

**Authors:** Zachary J.C. Tobias, Carolyn K. Tepolt

## Abstract

The repeatability of the genomic basis of adaptation depends on both genetic architecture and demographic history. Understanding this repeatability, and therefore the predictability of evolution, is increasingly important under global environmental change. Biological invasions are themselves a major component of global change and provide natural experiments for testing rapid adaptation and genomic parallelism across independently invaded ranges. The colonial ascidian *Botryllus schlosseri* is a widespread marine invader characterized by restricted larval dispersal and strong population structure. We used low-coverage whole-genome sequencing of 532 *B. schlosseri* individuals from 24 populations across North America to test for repeated thermal adaptation across the Atlantic and Pacific coasts. Genotype-environment association analyses identified thousands of loci associated with temperature in divergent lineages on either coast. Although most significant SNPs, genomic windows, and genes were lineage-specific, cross-lineage overlap exceeded null expectations, revealing a modest but detectable repeated component of putative thermal adaptation. Temperature-associated variation was broadly distributed across the genome. Regions with distinctive local population structure, potentially reflecting linked haplotypic structure or reduced recombination, showed modest enrichment for genotype-environment associations, particularly in the Pacific lineage, but did not disproportionately contain the parallel component of the response. Together, these results suggest that rapid thermal adaptation in *B. schlosseri* is partially repeatable despite deep divergence, but is largely distributed across the genome rather than concentrated within a few tightly linked regions. Restricted migration may therefore permit locally adaptive variants to evolve more independently across the genome, reducing the importance of concentrated genomic architectures favored under stronger migration-selection conflict.

## INTRODUCTION

The independent evolution of similar phenotypes in disparate lineages has long fascinated evolutionary biologists, both as compelling evidence for natural selection and for the fundamental questions it raises about evolutionary repeatability. Studies of repeated adaptation in natural populations and laboratory experiments have revealed outcomes ranging from reuse of the same mutations or genes to convergence at broader functional levels despite largely distinct underlying genetic changes (Barrick et al., 2009; Losos, 2011; Stern & Orgogozo, 2009; Wood et al., 2005). Accordingly, parallel evolution should be viewed as a continuum rather than a binary outcome, with the degree of repeatability potentially differing across biological scales from nucleotide variants to genes to pathways and ultimately phenotypes (Bolnick et al., 2018). Understanding what determines this variation in evolutionary repeatability has become especially relevant as populations increasingly encounter similar selective pressures as global environmental change accelerates.

The molecular repeatability of adaptation can be shaped by both the genetic architecture of selected traits and the demographic histories of evolving populations (Barrett & Schluter, 2008; Stern, 2013; Yeaman et al., 2018). These factors interact: demographic history influences which variants are available to selection, while genetic architecture can determine how effectively those variants respond under different demographic scenarios. Gene flow provides a particularly clear example. By introducing potentially maladaptive alleles and allowing recombination to break apart locally favored combinations, migration can impede adaptive divergence. Theory therefore predicts that adaptation under substantial migration can favor the concentration of adaptive variants in fewer, larger-effect regions, including genomic rearrangements that suppress recombination among locally favored loci (Tigano & Friesen, 2016; Yeaman, 2013; Yeaman et al., 2018). Empirical work also supports a connection between gene flow and the genomic architecture of adaptation, with adaptive differentiation under high connectivity often concentrated within tightly linked genomic regions or structural variants (Akopyan et al., 2026; Battlay et al., 2023, 2025; Shi et al., 2023). For example, across six riverine fishes spanning a gradient of population connectivity, putatively adaptive loci became increasingly clustered as gene flow increased (Shi et al., 2023). These patterns suggest that reduced-recombination regions can facilitate both local adaptation and evolutionary repeatability. Much theoretical work has therefore focused on how linkage and reduced recombination can facilitate local adaptation in the face of gene flow by limiting the breakup of locally favored combinations of alleles (Kirkpatrick & Barton, 2006; Yeaman, 2013; Yeaman & Whitlock, 2011). While theory provides clear expectations, we are still only beginning to understand how interactions between gene flow, genomic architecture, and repeated selection shape evolutionary responses in natural populations. Although large-effect loci and structural variants can contribute disproportionately to adaptation in some systems, distributed polygenic variation can also make substantial contributions (Koch et al., 2022; Stuart et al., 2023; Yeaman, 2015). Additionally, empirical approaches are often biased toward detecting large-effect loci and regions of reduced recombination, potentially underestimating the contribution of many individually small-effect variants (Hoban et al., 2016). *Botryllus schlosseri* provides a complementary case near the low-migration end of this continuum, offering an opportunity to test whether adaptation under limited gene flow is correspondingly less concentrated within tightly linked or recombination-suppressed genomic regions.

Biological invasions provide a particularly useful natural system in which to investigate this interaction between evolutionary repeatability, genomic architecture, and demographic history. Introduced populations routinely encounter novel environments and biological interactions, and local adaptation can arise over contemporary timescales (Lee, 2002; Prentis et al., 2008; van Boheemen et al., 2019). Independent introductions or range expansions can therefore act as natural evolutionary replicates, especially when genetically distinct populations encounter comparable environmental gradients. At the same time, replicate invasions may begin with different pools of genetic variation. Introductions can originate from distinct source populations, undergo demographic bottlenecks that stochastically reduce variation, or become admixed through interbreeding among repeated introductions from multiple sources (Bock et al., 2015; Kolbe et al., 2007; Rius & Darling, 2014; van Boheemen et al., 2017). These differences can push adaptation toward distinct molecular solutions even under similar selective pressures, whereas repeated use of the same variants, genomic regions, or pathways points to especially parallel components of the response. Studies of *Ambrosia artemisiifolia* have illustrated both outcomes, with repeated climatic adaptation across distinct invasion histories drawing in part on shared standing haploblocks (Battlay et al., 2023; Hodgins & Rieseberg, 2011; van Boheemen et al., 2019). Invasions therefore provide natural replicates for asking how differences in standing genetic variation and lineage history shape the degree and genomic basis of repeated adaptation.

The colonial ascidian *Botryllus schlosseri* provides a compelling system for exploring these ideas because it combines broad geographic spread with exceptionally restricted larval dispersal. *B. schlosseri* is a common member of marine fouling communities and has been transported widely among temperate coastlines, primarily through anthropogenic movement of adult colonies attached to vessels, aquaculture gear, or other mobile marine infrastructure (Carver et al., 2006; Lambert, 2001). In North America it occurs across broad latitudinal gradients on both the Atlantic and Pacific coasts. In the Pacific, *B. schlosseri* was first documented in the 1940s in San Francisco Bay (Carlton, 1979) but it has since spread between Sinaloa, Mexico in the south (Tovar-Hernández et al., 2014) and Sitka, Alaska in the north (Jurgens et al., 2018; Ruiz et al., 2006). Its invasive status is less certain in the Atlantic: it is considered cryptogenic along the mid-Atlantic and New England coasts of the United States but is definitely invasive in the Atlantic Maritime provinces of Canada, where its recent spread is well-documented (Carver et al., 2006; Fofonoff et al., 2024; Yund et al., 2015). Earlier mitochondrial and microsatellite studies have shown that North American populations are strongly structured and that Atlantic and Pacific populations derive from genetically distinct sources rather than a single recent transcontinental expansion (Bock et al., 2012; Lacoursière-Roussel et al., 2012; Lejeusne et al., 2011; Stoner et al., 2002). More broadly, multilocus surveys have revealed deep phylogeographic structure within the *B. schlosseri* species complex, with the globally widespread clade A occupying a much broader range than several more geographically restricted lineages in Europe (Bock et al., 2012; Reem et al., 2021). As taxonomic characterization of these lineages is ongoing (Brunetti et al., 2020), we use *B. schlosseri* to refer solely to the globally widespread clade A, which is the species present in North America.

Natural dispersal in *B. schlosseri* occurs on an extraordinarily small spatial scale. Direct estimates indicate that most larvae settle within approximately one meter of their birthplace (Grosberg, 1987), and genetic differentiation has been detected over tens of meters within continuous populations (Yund & O’Neil, 2000). Population-genetic studies using microsatellites and mitochondrial markers likewise reveal strong differentiation among sites, temporal turnover, and substantial local structure across introduced populations (Lin & Zhan, 2016; Reem et al., 2013; Stoner et al., 2002). Despite its extensive range expansion, *Botryllus schlosseri* lacks the persistent, homogenizing migration characteristic of many marine species with longer-lived planktonic larvae. This demographic regime may be especially conducive to rapid and locally idiosyncratic adaptation. Once a population is established, locally favorable alleles can potentially increase without being continually broken apart or swamped by immigrants. At the same time, tunicates are notable for high levels of nucleotide polymorphism, rapid sequence evolution, and elevated recombination rates (Berna & Alvarez-Valin, 2014; Denoeud et al., 2010; Kano et al., 2006; Tsagkogeorga et al., 2012). Together, abundant genetic variation, strong subdivision, and limited larval exchange provide a distinctive genomic and demographic substrate for local adaptation.

Temperature is a particularly relevant axis of selection in this and many other marine systems, as climate change alters thermal environments across species’ ranges and exposes geographically separated populations to similar selective pressures (Hoffmann & Sgró, 2011; McGaughran et al., 2021). Across North America, *B. schlosseri* spans pronounced latitudinal gradients in coastal temperature, including recently colonized northern environments. Previous physiological studies found latitudinal differentiation in heat and cold tolerance across populations on both coasts, consistent with the potential for repeated thermal adaptation (Tobias et al., 2024, 2025). The independent Atlantic and Pacific ranges therefore provide an opportunity to test whether similar thermal gradients produce repeated genomic responses from strongly divergent genetic backgrounds, and whether those responses have the same genomic architecture seen in systems with higher gene flow.

Here, we use low-coverage whole-genome sequencing of 532 *B. schlosseri* individuals from 24 populations spanning both coasts of North America, to investigate the genomic architecture and repeatability of putative adaptation to environmental temperature. Population genomic analyses identify two widespread, highly differentiated lineages that independently span broad thermal gradients and therefore provide natural evolutionary replicates. Using these lineages, we ask three questions. First, does genomic variation track environmental temperature within each lineage, suggestive of local thermal adaptation? Second, to what extent are temperature-associated signals repeated between the two lineages at the level of individual SNPs, genomic windows, and genes, and what functions are represented among shared candidate genes? Third, is putatively adaptive variation concentrated in extended genomic regions with distinctive local population structure, as might arise from reduced recombination or structural variation, or is it more diffusely distributed across the genome? By addressing these questions in a species with exceptionally restricted routine dispersal, we test how demographic context shapes both the repeatability and genomic architecture of adaptation to environmental change.

## MATERIALS AND METHODS

### Collections and DNA Extractions

We collected 25-40 *B. schlosseri* colonies from each of 24 sites along the Atlantic and Pacific coasts of North America between August 2019 and September 2022 (Figure 1). Collections were spaced by at least two meters when possible to reduce sampling of close kin. Colony fragments were preserved in 95% ethanol and stored at -80 °C. After removing the tunic, gut, and brooding embryos, DNA was extracted from tissue pooled from zooids from the same or adjacent systems of each colony using Qiagen DNeasy Blood and Tissue kits. Additional collection and extraction details are provided in the Supporting Information.

**Figure 1.**
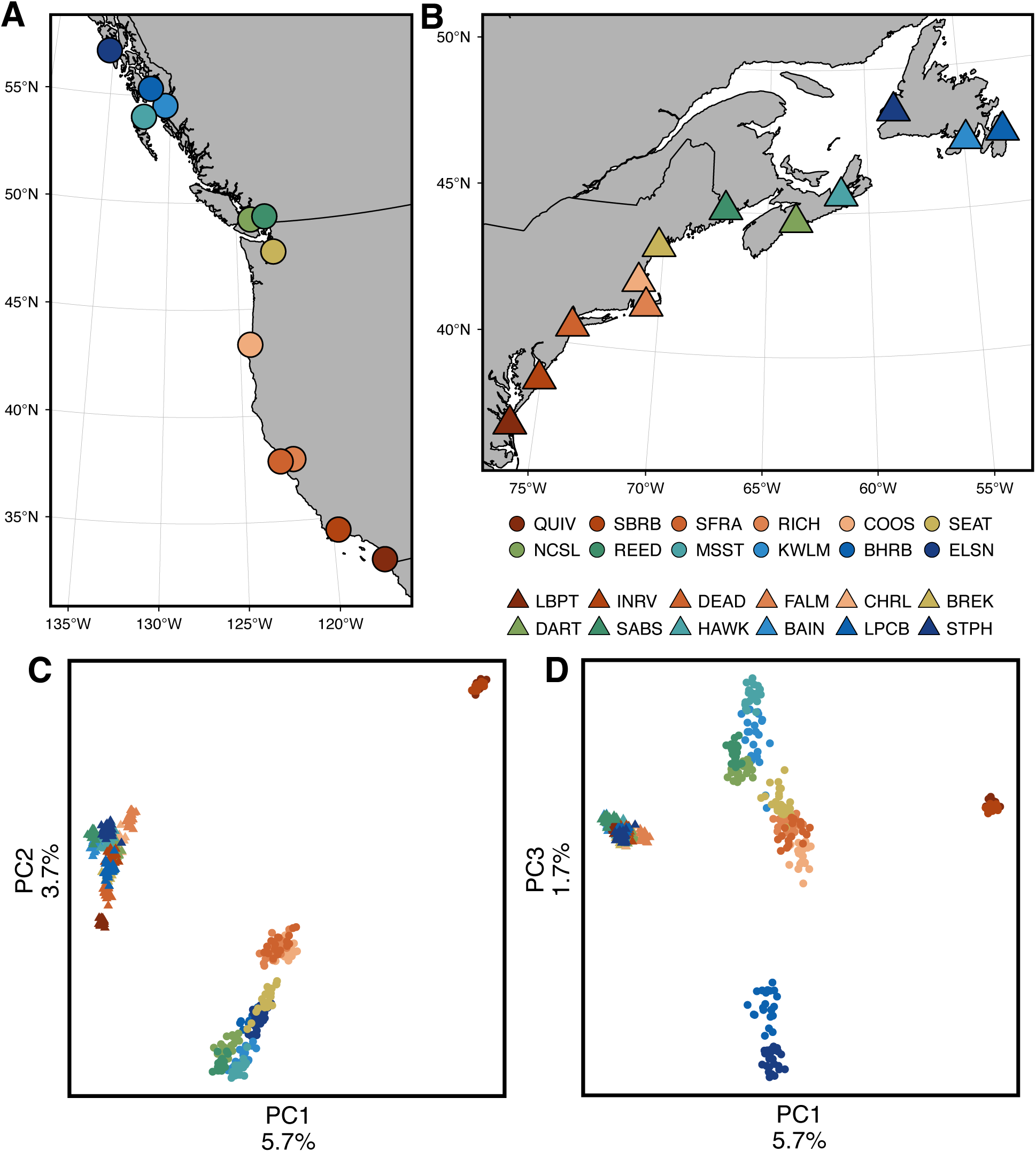
Sampling and population genetic structure. A-B) Sampling locations of *B. schlosseri* along Pacific (A) and Atlantic (B) coasts of North America. C-D) Principal components analyses (PCAs) of genotype likelihoods across 151,289 LD-pruned SNPs. Clustering patterns reveal three broad lineages of *B. schlosseri* in C): Northwest Atlantic (NWA) in left-center, Northeast Pacific (NEP) in bottom-center, and Southern California (SoCal) in top-right. Individuals from Southeast Alaska (BHRB and ELSN) separate from the NEP cluster along PC3 (D).

### Library preparation and sequencing

Sequencing libraries were prepared from 540 specimens following modified versions of Baym et al. (2015) and Therkildsen & Palumbi (2017). Briefly, 1-10 ng of genomic DNA was tagmented, amplified to add Illumina adapters and unique dual indexes, pooled, and electrophoretically size-selected for fragments of 500–8,000 bp. Equimolar pools were sequenced across six lanes of three NovaSeq 6000 SP flow cells using 150 bp paired-end reads at the University of Utah High-throughput Genomics Center. Complete library-preparation details are provided in the Supporting Information.

### Bioinformatic processing

The bioinformatic pipeline was adapted from lcWGS workflows described by Therkildsen & Palumbi (2017), Lou et al. (2021), and Zhou et al. (2024), with full parameters provided in the Supporting Information. One of 540 libraries produced no assigned reads. Reads from the remaining libraries were evaluated with FastQC v0.12.1 (Andrews, 2010) and MultiQC v1.27.1 (Ewels et al., 2016), trimmed with Trim Galore v0.6.10 (Krueger, 2015; Martin, 2011), and aligned to the *B. schlosseri* reference genome (De Thier et al., 2025) using Bowtie2 v2.5.4 (Langmead & Salzberg, 2012) in -- very-sensitive-local mode. One specimen with extremely low mapping was identified from mitochondrial sequence as *Botrylloides violaceus* and removed.

BAM files were filtered for mapping quality and concordant pairs. Overlapping read segments were soft-clipped with bamUtil v1.0.15 (Jun et al., 2015), PCR duplicates were removed with Picard v2.26.3 MarkDuplicates (“Picard Toolkit,” 2018), and reads were realigned around indels with GATK v3.8.1 (McKenna et al., 2010).

We used ANGSD v0.938 (Korneliussen et al., 2014) to estimate genotype likelihoods and allele frequencies. After an initial coverage and missingness screen, six additional low-quality samples were removed. ANGSD was rerun by chromosome using the SAMtools likelihood model (Danecek et al., 2021), requiring data from at least half of the retained individuals and imposing an empirical upper-depth threshold to reduce repetitive or paralogous signal. Four individuals from Dartmouth, NS (DART) were identified as probable migrants or mislabeled samples were excluded from analyses requiring population assignment, but retained in lineage-wide analyses such as local PCA.

### Population structure and linkage disequilibrium

To characterize broad population structure, we filtered genotype-likelihood data by minor allele frequency and missingness, thinned sites before LD estimation, and used ngsLD v1.2.0 (Fox et al., 2019) and its prune_graph utility to generate approximately unlinked SNP sets. These pruned genotype likelihoods were analyzed with PCAngsd v1.03 (Meisner & Albrechtsen, 2018). PCA results were used to identify major population clusters and finer subclusters. We then repeated ANGSD, LD pruning, and PCAngsd analyses within clusters to refine population assignments. Pairwise population F_ST_ was calculated by estimating folded two-dimensional site-frequency spectra with winSFS v0.7.0 (Rasmussen et al., 2022) and then applying realSFS fst index and fst stats; full site filtering and SFS procedures are provided in the Supporting Information.

To characterize the physical scale of linkage disequilibrium independently of broad among-population structure, we additionally estimated LD within each of the 22 focal NWA and NEP populations, excluding the two Southern California populations, using ngsLD. Sites with estimated minor allele frequency of at least 0.05 were pre-thinned to a minimum separation of 100 bp for computational efficiency, and genotype-likelihood-based LD was calculated for pairs separated by no more than 50 kb. We summarized r2_ExpG in 100-bp distance bins and defined the characteristic decay distance as the first of ten consecutive bins for which mean r2_ExpG remained at or below 0.2. Requiring ten consecutive bins reduced sensitivity to noise in individual bins and made this a conservative estimate of the distance over which appreciable LD persisted. We used binned empirical summaries rather than nonlinear decay curves because curve fitting was unstable for these low-coverage data.

### Nucleotide diversity

We estimated nucleotide diversity within each population from genotype likelihoods at a common set of 204,842,689 callable sites (both fixed and variant). Folded one-dimensional site-frequency spectra estimated with winSFS were supplied to ANGSD/realSFS saf2theta. Genome-wide π was calculated as the sum of pairwise theta (tP) across the 16 chromosomes divided by the summed number of analyzed sites. We tested its relationship with sample size using Spearman rank correlation. Population-level estimates are provided in the accompanying source data.

### In situ temperature data

Environmental temperature data were derived from publicly available *in situ* instrumentation (shore-deployed sondes, ocean-moored buoys, etc.; Table S1). We primarily used data from January 1st, 2015 through October 19th, 2023, though data availability and quality constrained this range at some sites. We initially extracted a variety of temperature metrics including the mean number of days per year exceeding 12 °C (the approximate threshold of reproduction for *B. schlosseri*), mean annual temperature, mean winter temperature, and mean summer temperature. We also included site latitude as a potential explanatory factor. Because these were all highly collinear, we retained mean annual temperature as the sole environmental covariate for BayPass v2.1 analyses (Gautier, 2015).

### Genotype-environment association analyses

For genotype-environment association analyses, we focused on the two most widespread population clusters identified by PCA: the Northwest Atlantic (NWA) and Northeast Pacific lineages (NEP) (see Results). The two southern California populations formed a distinct subcluster and were not used in downstream GEA analyses. ANGSD was run separately for each population to estimate allele frequencies. We retained sites with data from at least 10 of 12 NWA populations or 9 of 10 NEP populations and a lineage-wide minor allele frequency of at least 0.05. This yielded final BayPass input matrices containing 8,770,990 SNPs for NWA and 8,858,634 SNPs for NEP.

Within each lineage, we estimated a neutral covariance matrix (omega matrix) from an LD-pruned subset of SNPs using the BayPass neutral model. We then used a BayPass genotype-environment association model (--covmcmc) to test for associations between allele frequencies and mean annual temperature separately within each lineage.

### Investigating parallel GEA signals

Because BayPass GEA scores were highly consistent across five random seed replicates, we summarized each SNP using the median eBPmc across replicates. SNPs with median eBPmc ≥ 3, corresponding to an empirical Bayesian p-value < 0.001, were treated as significant. We examined parallelism at the level of individual loci, genomic windows, and genes.

At the SNP level, we tested whether temperature-associated allele-frequency effects were directionally concordant across lineages. For each focal lineage, we retained SNPs with eBPmc ≥ 3, assigned them to 50 kb intervals, and repeatedly sampled one SNP per occupied interval. This 50 kb thinning scale was deliberately broad so that a permutation replicate did not repeatedly draw nearby signals likely to be non-independent because of linkage or other sources of spatial autocorrelation. For each resample, we calculated two concordance statistics: the proportion of SNPs with the same effect-size (M_Beta) direction across lineages, and the mean product of cross-lineage M_Beta values. Null distributions were generated by randomly flipping the sign of one lineage’s M_Beta values, thus preserving the SNP set, spatial thinning, and the distribution of absolute effect sizes while randomizing cross-lineage sign alignment. For this test and the window- and gene-level permutation tests described below, we performed 1,000 iterations. One-sided empirical p-values were calculated as [1 + the number of null statistics greater than or equal to the observed statistic]/1,001. The cross-lineage M_Beta product was our direct magnitude-weighted statistic. Loci with same-sign but near-zero effects contribute little to this statistic, whereas opposite-sign effects contribute negatively. It therefore tests whether concordance is concentrated among loci with larger effects without imposing an arbitrary absolute-M_Beta threshold.

For window-level analyses, we tiled the genome into non-overlapping windows and summarized BayPass signal separately within each lineage. The focal analysis used 5 kb windows, with 500 bp and 50 kb windows used as sensitivity analyses for narrower and broader genomic signals, respectively. For each window, we recorded the number of BayPass SNPs, the maximum eBPmc, and the mean eBPmc in each lineage. We used two complementary definitions of window-level signal. The first classified windows containing at least one significant SNP (maximum eBPmc ≥ 3), capturing a strong association even if represented by a single analyzed SNP. The second classified windows in the top 5% of mean eBPmc within each lineage, capturing more broadly elevated evidence across the SNPs in a window though potentially below the eBPmc ≥ 3 threshold. A window was considered parallel if it met the relevant signal definition in both NWA and NEP.

To test whether the observed number of parallel windows exceeded expectation, we used matched and circular null models. Both were restricted to windows containing at least one BayPass SNP in each lineage. The matched null controlled for differences in the opportunity to detect associations by permuting windows within chromosomes and SNP-density bins, defined from deciles of SNP number in each lineage. Within each matched bin, we permuted the NEP statistic relative to NWA and recalculated the number of parallel windows. One-sided empirical p-values were calculated as above.

The chromosome-wise circular permutation retained broad-scale genomic clustering of GEA signal. Significant or elevated windows from one lineage were shifted by a random distance within each chromosome, wrapping around chromosome ends, and then overlapped with windows from the other lineage. This preserved the number and spatial clustering of outlier windows within chromosomes while testing whether their cross-lineage alignment was greater than expected. The matched and circular tests therefore address complementary features of the null expectation: heterogeneous SNP density and broad spatial non-independence, respectively.

For gene-level analyses, we identified genes containing BayPass SNPs in their exonic sequence. We used two definitions of gene-level signal that mirrored the window-level analyses. First, a gene was considered significant in a lineage if at least one significant BayPass SNP in that lineage overlapped one of its exons. Second, for genes with exonic BayPass SNPs, we calculated the mean eBPmc across all exonic BayPass SNPs in each lineage and classified genes in the top 5% of mean exonic eBPmc values as elevated. Genes were considered parallel candidate genes if they met the relevant signal definition in both NWA and NEP.

For gene-level null tests, we used matched permutations to account for differences in opportunity among genes. We restricted the background to genes with at least one exonic BayPass SNP in both lineages. For the significant-SNP definition, genes were permuted within exon-length bins, using deciles of total reduced exon length. For the mean-eBPmc definition, genes were permuted within bins defined by the number of exonic BayPass SNPs, using deciles of exonic SNP count. Within each matched bin, we randomly sampled the observed number of NWA-hit and NEP-hit genes and recalculated the overlap between the two sampled sets. This overlap was used as the null number of parallel genes.

### Functional annotation and enrichment

Predicted proteins were annotated with eggNOG-mapper v2.1.13 (Cantalapiedra et al., 2021) against eggNOG v5.0.2 (Huerta-Cepas et al., 2019) using DIAMOND v2.0.15 (Buchfink et al., 2015). We retained orthology-based annotations with e-values < 1 × 10-10. The conservative candidate list comprised genes containing significant exonic BayPass SNPs in both lineages. For functional enrichment, we used the broader set of genes containing significant SNPs anywhere in the gene body, thereby including putatively regulatory variation.

We used clusterProfiler (Yu et al., 2012) to test GO-term over-representation against the background of genes containing at least one analyzed SNP in both lineages. Redundant significant terms across all three GO namespaces were clustered by Jaccard similarity of member genes, and clusters were labeled from their constituent terms and gene annotations.

### Extended regions of distinctive local population structure

To identify extended genomic regions whose patterns of local population structure differed from the genome-wide background, we used the R package lostruct v0.0.0.9000 (Li & Ralph, 2019). For each lineage separately, we used PCAngsd to estimate sample covariance matrices in non-overlapping 100 kb windows and retained windows containing at least 250 SNPs after PCAngsd filtering. lostruct compares the geometry of local PCA representations among windows; we used the first two local PCs to calculate inter-window distances, performed multidimensional scaling on the resulting distance matrix, and retained the first five MDS axes. This procedure detects genomic heterogeneity in local relatedness or covariance patterns. Such heterogeneity can arise from inversions or reduced recombination, but also from linked selection, recombination-rate variation, admixture, or demographic history, so lostruct alone does not identify a structural variant or its mechanism. To summarize the degree to which each 100 kb window departed from the genome-wide pattern of local population structure, we calculated its Euclidean distance from the genome-wide median position across the first five MDS axes and ranked these distances within each lineage.

We also quantified how strongly local genetic variation was structured within each 100 kb window using the total proportion of variance explained by PC1 and PC2 from the corresponding PCAngsd covariance analysis. For a complementary categorical analysis, we classified windows as having distinctive, dominant-axis local population structure when they fell in the top 10% for both Euclidean MDS distance and total variance explained by PC1 + PC2 within a lineage. This joint criterion identified windows that were both unusually distant from the genome-wide pattern and dominated by a strong local axis of genetic differentiation. Calls were made and tested at the individual-window level; contiguous qualifying windows were combined only for descriptive summaries and no intervening gaps were bridged.

### Association of local genomic structure with GEA and LD

To test whether genotype-environment association signal was disproportionately concentrated in regions with distinctive local population structure, we summarized significant BayPass signal on the same 100 kb scale used for local PCA. We tested associations with both the continuous lostruct score and the categorical set of local-structure candidate windows, considering lineage-specific and parallel GEA signal. Significance was assessed using chromosome-wise circular permutations to account for spatial autocorrelation along chromosomes. Full details are provided in the Supporting Information.

We also asked whether regions with distinctive local population structure exhibited elevated within-population LD. Population-specific ngsLD estimates for SNP pairs separated by 5–50 kb were summarized within each 100 kb local-PCA window and standardized within populations and chromosomes before being compared with lostruct scores. Associations were tested using Spearman’s rank correlation with chromosome-wise circular permutations. Full details are provided in the Supporting Information.

### Exploratory characterization of the chromosome 6 region

We further characterized the prominent NEP chromosome 6 region using local PCA across the focal interval and adjacent regions. We also compared PCAngsd inbreeding coefficients among the three observed local-PCA classes to ask whether they exhibited the heterozygosity pattern expected under a simple two-arrangement inversion model. Because these analyses are not diagnostic of structural variation, we treat the region as an illustrative example of distinctive local genomic structure rather than a confirmed inversion.

## RESULTS

### Population genetic structure and genomic diversity

We generated low-coverage whole-genome sequence data for 538 *Botryllus schlosseri* individuals from 24 populations across its range on both coasts of North America (Figure 1A-B). After dropping low-depth outliers, we retained 532 individuals with a median whole-genome depth of 0.79× (IQR: 0.70-0.90×; Supplementary Figure S1). For a first analysis comprising all individuals, we obtained genotype likelihoods at 14,021,658 SNPs with estimated minor allele frequencies ≥ 0.05. PCA on an LD-pruned subset of 151,289 SNPs revealed three distinct clusters of populations: one containing all samples from the Northwest Atlantic (NWA), a second containing Northeast Pacific samples from San Francisco Bay northward (NEP), and a third representing two populations from southern California (SoCal) (Figure 1C-D). To ensure this clustering was not driven by asymmetric patterns of site missingness across populations or clusters, we repeated PCA using sites covered in at least 90% of individuals and obtained similar results (Supplementary Figure S2). We re-estimated genotype likelihoods within each cluster to identify potential migrants and examine finer patterns of population structure. This revealed hierarchical population structure and strong clustering by population (Supplementary Figure S3), though these patterns will be treated more comprehensively elsewhere (Tobias and Tepolt, in prep.).

Estimates of pairwise F_ST_ among populations indicated substantial genetic differentiation both across and within clusters (Figure 2A-B). The SoCal cluster exhibited very strong differentiation from both the NEP and NWA clusters (mean pairwise FST of 0.32 and 0.40, respectively). Mean pairwise FST between NEP and NWA populations was 0.19 (range: 0.13-0.27). Mean pairwise FST was 0.10 (range: 0.03–0.18) among NEP populations and 0.11 (range: 0.05-0.22) among NWA populations.

**Figure 2.**
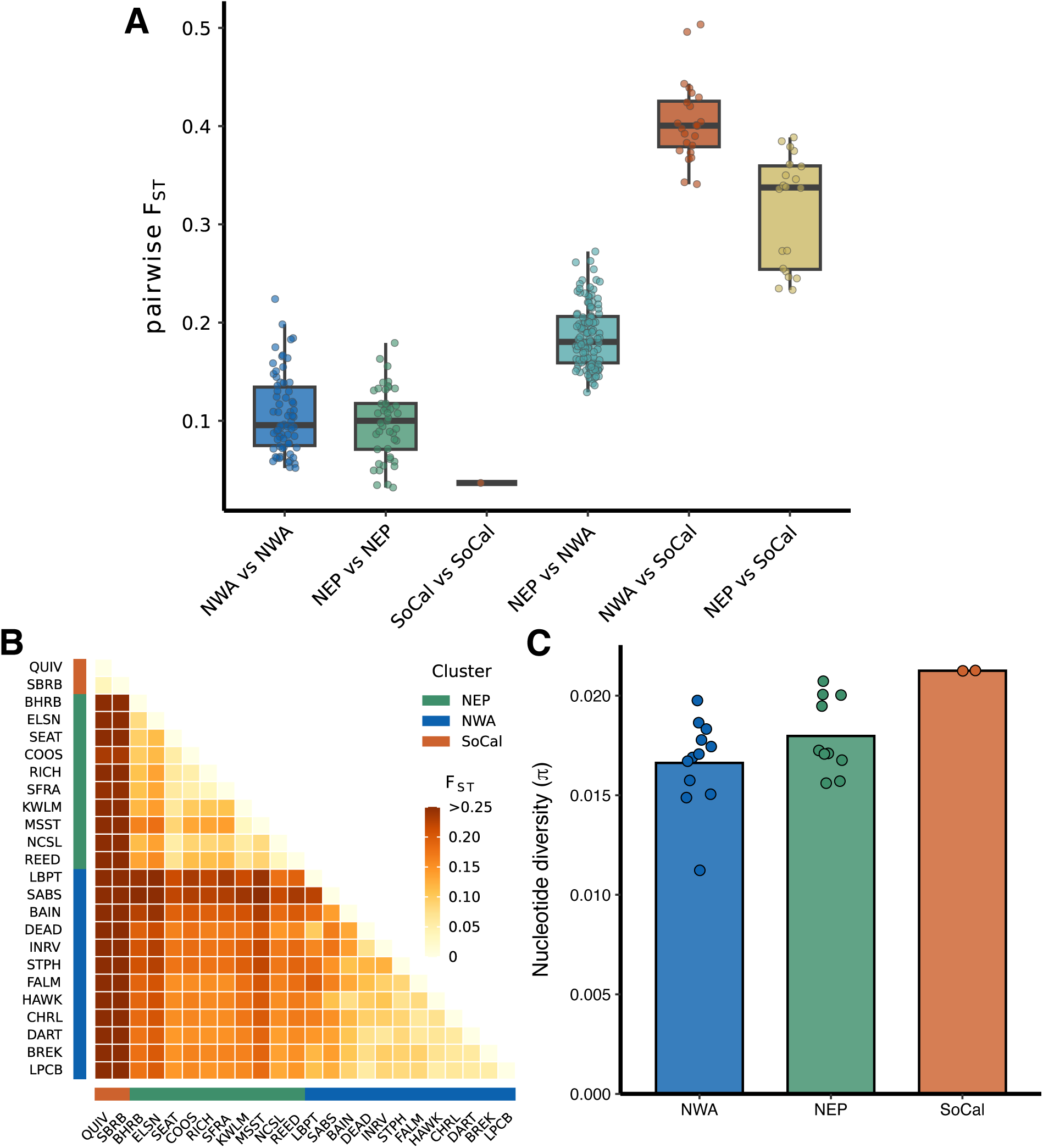
Strong within- and among-lineage divergence in *B. schlosseri*. A) Box plots of pairwise-F_ST_ of populations within and across lineages identified from PCA. Points represent individual population pairs. B) Heatmap of pairwise F_ST_ among all population pairs. Populations are ordered by hierarchical clustering, not lineage assignment. C) Bar plot of mean nucleotide diversity (π) estimates across lineages. Points represent individual population estimates.

Within-population linkage disequilibrium decayed rapidly with physical distance. Across the 22 focal NWA and NEP populations, mean genotype-likelihood-based LD declined below r2_ExpG = 0.2 and remained below that threshold by approximately 1.0–1.2 kb. Because this estimate was based on noisy low-coverage summaries and required ten consecutive bins below the threshold, it should be interpreted as a conservative upper bound on the characteristic decay scale; LD for many comparisons likely decayed sooner. We therefore treated 5 kb as the focal aggregation scale for GEA parallelism, with 500 bp and 50 kb representing finer- and broader-scale sensitivity analyses, respectively.

Nucleotide diversity was substantial within populations but varied geographically (Figure 2C; Supplementary Table S2). Genome-wide π ranged from 0.0156 to 0.0207 among NEP populations (mean = 0.0180; median = 0.0172) and from 0.0112 to 0.0198 among NWA populations (mean = 0.0166; median = 0.0170). The two SoCal populations had π = 0.0212–0.0213. Population π was unrelated to the number of sampled individuals (Spearman’s rho = 0.033, p = 0.877), indicating that the among-population differences were not an obvious consequence of the modest variation in sample size.

### Genotype-environment association analysis

The NEP and NWA lineages both have broad latitudinal ranges spanning steep thermal gradients (Supplementary Table S1). We used BayPass within each lineage to detect associations between population-level allele frequency and the mean annual temperature of each sampling location, as determined from publicly available *in situ* water temperature records. The final analyses contained 8,770,990 SNPs in NWA and 8,858,634 SNPs in NEP.

After accounting for underlying neutral covariance structure, we obtained 27,470 and 17,938 significantly associated SNPs (median eBPmc ≥ 3) for NEP and NWA, respectively. Significant temperature-associated SNPs were broadly distributed across the genome (Figure 3A-B), although the density of GEA signal varied among genomic regions. In NEP, this included one extended and sharply defined peak of elevated GEA signal near the start of chromosome 6 (Figure 3A), which we examine in greater detail below.

**Figure 3.**
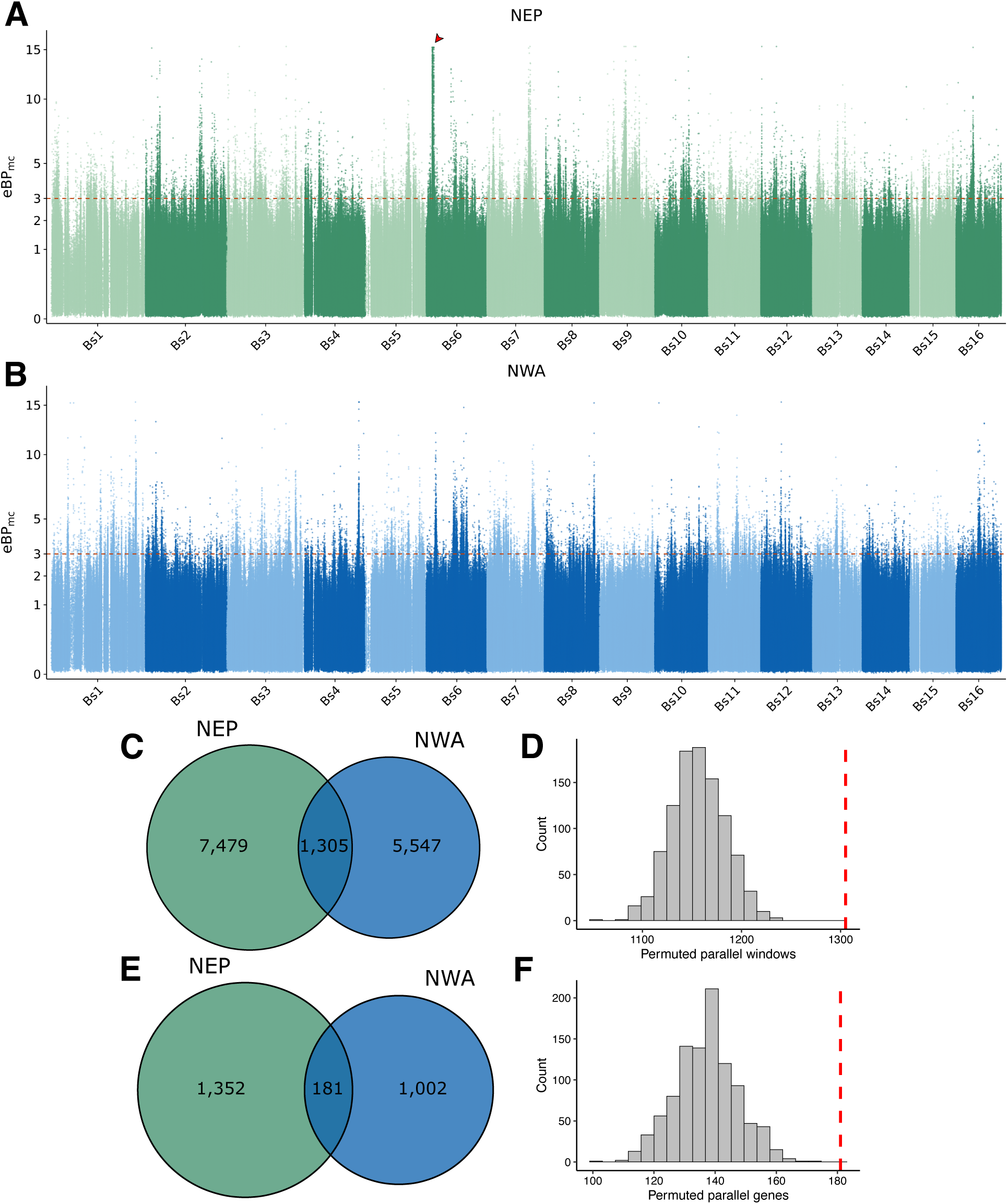
A diffuse but partially repeated architecture of thermal adaptation across the NEP and NWA lineages of *B. schlosseri*. A-B) Manhattan plots of Baypass eBP_mc_ (approximate empirical Bayesian p-values) across the genome for NEP (A) and NWA (B). The red arrow in A) represents a region of atypical local covariance on chromosome 6 described in the main text. C) Overlap of 5kb windows containing a significant BayPass SNP (eBP_mc_ ≥ 3) in NEP and NWA. D) Null distribution of parallel 5-kb windows under a SNP-density-matched permutation test; the red dashed line denotes the observed overlap. E) Overlap of genes containing a significant exonic BayPass SNP in NEP and NWA. F) Null distribution of parallel genes under an exon-length-matched permutation test.

### Parallelism of GEA signal

We investigated parallelism of GEA signal at the level of individual loci, genomic windows, and genes. Of the 27,470 and 17,938 temperature-associated SNPs in NEP and NWA, respectively, only 161 were significant in both lineages. Because SNPs were identified and filtered in each lineage separately and were therefore subject to ascertainment differences, we assessed SNP-level parallelism by testing whether loci shared across the two BayPass datasets exhibited concordance in the direction and magnitude of their association with temperature. A total of 5,325,358 loci were shared between the NEP and NWA analyses.

For NWA-ascertained temperature-associated SNPs, 52.8% of resampled SNPs had the same sign of temperature association in both lineages, compared with 50.1% under the sign-randomized null (p = 0.016). The magnitude-weighted statistic, calculated as the mean product of cross-lineage M_Beta values, was also elevated relative to the null (p = 0.007). For NEP-ascertained temperature-associated SNPs, 51.1% had concordant signs across lineages, which was not significantly different from the sign-randomized expectation (p = 0.170), whereas the M_Beta-product statistic was elevated (p = 0.033). Because same-sign loci with effects near zero contribute little to this product and opposite-sign loci contribute negatively, the magnitude-weighted tests show that concordance was concentrated disproportionately among loci with larger environmental-association effect estimates. Together, these results indicate weak but detectable site-level parallelism in temperature-associated allele-frequency shifts.

Next, we aggregated each lineage’s SNP-level GEA signal to 5 kb windows, with 500 bp and 50 kb windows used as sensitivity analyses. In NEP, 8,784 5 kb windows contained at least one SNP with eBPmc ≥ 3, whereas 6,852 such windows were identified in NWA. Of these, 1,305 windows were shared between NEP (14.9%) and NWA (19.1%) (Figure 3C). For the mean-eBPmc approach, we took the upper 5% of windows in each lineage (NEP: 3,125; NWA: 3,144) and found that 193 windows were shared.

We conducted permutations to test whether these overlaps were greater than expected. For the maximum-eBPmc approach, we observed 13% more shared 5 kb windows than expected under the chromosome- and SNP-density-matched null (observed: 1,305; null mean: 1,156.3; one-sided p = 0.001) (Figure 3D). Under the chromosome-wise circular null, which additionally preserved broad spatial clustering, we observed 48% more overlapping significant 5 kb windows than expected (null mean: 880.1; p = 0.001). The significant-window result also held at 500 bp under both nulls but not at 50 kb under the circular null. For the upper-5%-mean definition, overlap was supported at 500 bp and 5 kb under both nulls but not at 50 kb. Complete permutation summaries are provided in the accompanying source data.

We also examined the degree of GEA overlap at the gene level. Considering only exonic SNPs, we identified 1,533 genes containing at least one SNP with eBPmc ≥ 3 in NEP and 1,183 in NWA. Of these, 181 were shared between NEP (11.8%) and NWA (15.3%) (Figure 3E). A permutation test matching genes within exon-length deciles revealed significantly greater overlap than expected (null mean: 137.2; enrichment = 1.32; one-sided p = 0.001) (Figure 3F). Overlap among the top 5% of genes ranked by mean exonic eBPmc was also elevated (observed: 53; null mean: 40.7; enrichment = 1.30; p = 0.032).

### Parallel candidate genes and functional enrichment

Of the 181 genes containing at least one significant exonic BayPass SNP in both lineages, 113 were assigned to an orthologous group by eggNOG-mapper with an e-value ≤ 1 × 10-10 (Supplementary Table S3). These included orthologs involved in mitochondrial respiration, redox homeostasis, and cellular stress responses, including succinate dehydrogenase complex iron-sulfur subunit B (SDHB), ferritin heavy chain (FTH1), glutathione S-transferase kappa 1 (GSTK1), and eukaryotic translation initiation factor 2-alpha kinase 3 (EIF2AK3). Other candidates were associated with environmental sensing, growth and metabolic regulation, and retinoic acid metabolism, including aryl hydrocarbon receptor (AHR), serine/threonine-protein kinase TOR (MTOR), retinol dehydrogenase 10 (RDH10), and cytochrome P450 family 26 subfamily A member 1 (CYP26A1).

To explore functional enrichment, we conducted over-representation analysis of GO terms using a broader set of parallel genes, defined as genes containing significant BayPass SNPs anywhere in the gene body. This set comprised 955 genes. After clustering redundant significant GO terms (Supplementary Table S4) based on the Jaccard overlap of constituent genes, we identified nine broad functional categories enriched for parallel genes (Figure 4). These included functions related to immune and endocrine signaling, neurovascular morphogenesis, and cell adhesion, among others.

**Figure 4.**
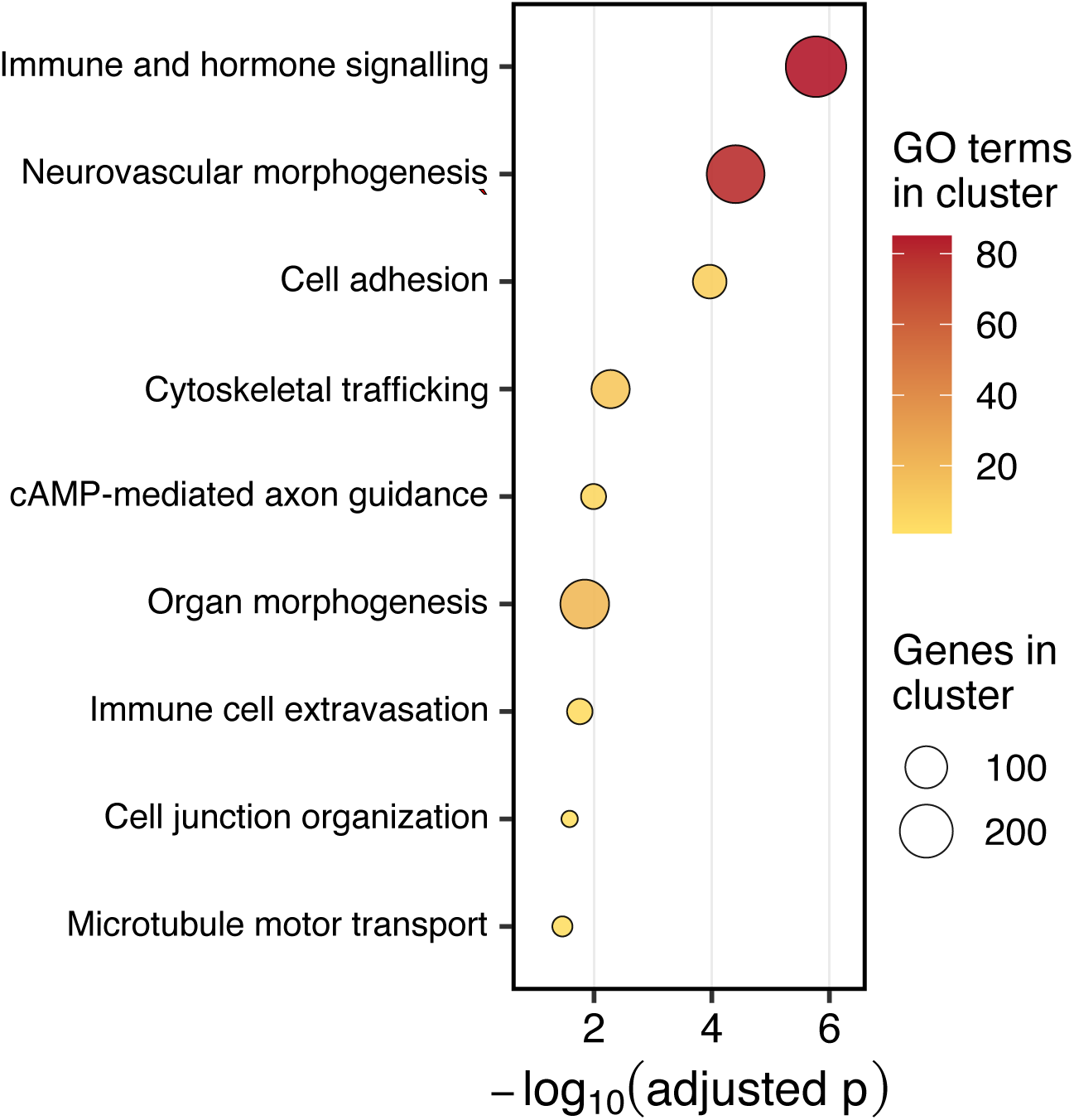
Functional enrichment among parallel candidate genes. Gene Ontology (GO) enrichment was performed using genes containing significant BayPass SNPs anywhere in the gene body in both NWA and NEP. Significant GO terms across all three GO namespaces were clustered according to Jaccard similarity in their constituent gene sets to reduce redundancy, and each cluster was assigned a representative functional label through manual interpretation of the contributing GO terms and genes.

### GEA is weakly associated with distinctive local population structure, more so in NEP

Local PCA revealed substantial heterogeneity in population structure across the genome in both lineages. The Euclidean distance from the genome-wide median position in MDS1-MDS5 provided a continuous lostruct score for each 100 kb window, while the proportion of variance explained by local PC1 + PC2 quantified the strength of the dominant local axes. Genome-wide score tracks and categorical calls are shown in the Supporting Information, including their spatial clustering along chromosomes.

GEA signal generally increased with the lostruct score in both lineages (Figure 5A). The proportion of 5 kb windows containing a significant BayPass SNP was weakly positively correlated with the lineage-specific score percentile in NWA (Spearman’s rho = 0.139, circular-permutation p = 0.001) and more strongly, although still modestly, in NEP (rho = 0.245, p = 0.001). Parallel GEA was also weakly associated with the larger of the two lineage-specific score percentiles (rho = 0.094, p = 0.001). Thus, GEA was not distributed independently of local population structure, but the associations were modest and strongest in NEP.

**Figure 5.**
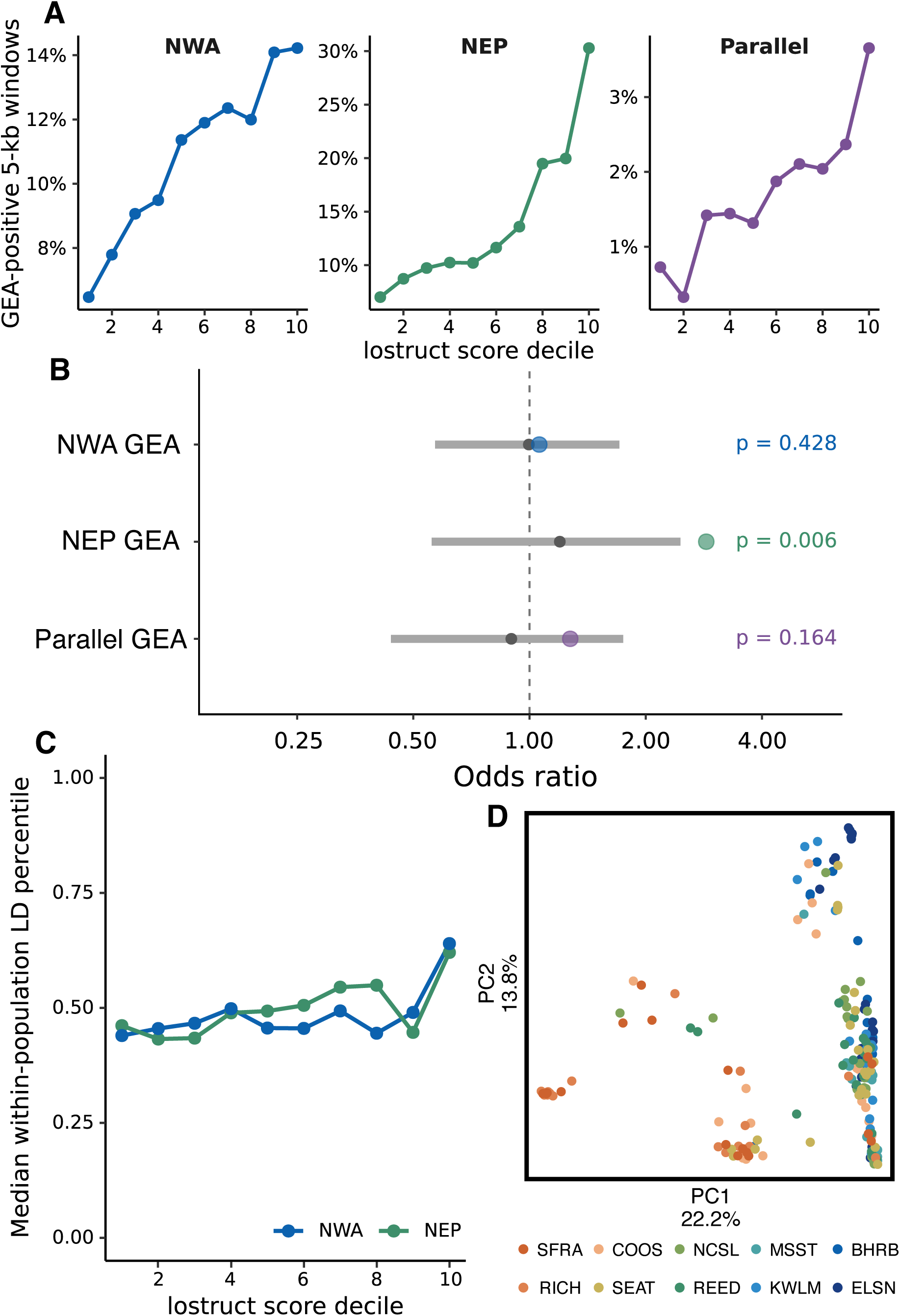
Genotype-environment associations and linkage disequilibrium in regions of distinctive local population structure. (A) Proportion of 5-kb windows containing significant GEA signal across deciles of the 100-kb lostruct score for NWA, NEP, and parallel GEA. Parallel windows contained significant GEA signal in both lineages. (B) Enrichment of GEA within candidate local-structure windows, defined as 100-kb windows in the top 10% for both lostruct score and variance explained by local PC1 + PC2. Colored points show observed odds ratios; gray points and intervals show the corresponding circular-permutation null distributions. Empirical one-sided p-values are shown. For parallel GEA, candidate status was defined by meeting the local-structure criterion in either lineage. (C) Median within-population LD percentile for SNP pairs separated by 5-50 kb across lostruct score deciles in NWA and NEP. (D) Local PCA of the prominent NEP chromosome 6 interval (Bs6:3.145-3.950 Mb), colored by sampling population, illustrating the localized three-cluster population structure observed in this region.

The stricter categorical definition identified 112 of 4,392 NWA windows (2.55%) and 110 of 4,422 NEP windows (2.49%) that fell in the top 10% for both lostruct score and variance explained by PC1 + PC2. These candidate windows did not contain appreciably more NWA GEA than background windows (11.6% versus 11.0% of analyzable 5 kb windows; odds ratio = 1.06, circular-permutation p = 0.428). In NEP, however, 31.4% of 5 kb windows inside candidates contained GEA signal compared with 13.7% outside them (odds ratio = 2.87, p = 0.006; Figure 5B). This enrichment was not driven by the conspicuous chromosome 6 region: after removing all nine overlapping 100 kb windows, the NEP effect was essentially unchanged (odds ratio = 2.89, p = 0.006), as was the continuous relationship (rho = 0.243, p = 0.001).

Candidate status in either lineage was not significantly associated with parallel GEA (odds ratio = 1.27, p = 0.164; Figure 5B). Together, the continuous and categorical analyses show that temperature-associated variation tends to accumulate weakly in regions with unusual local population structure, particularly in NEP, but provide no evidence that the most extreme dominant-axis regions disproportionately contain the repeated component of GEA. The continuous association is also not independent evidence of reduced recombination: allele-frequency shifts produced by selection can themselves contribute to both BayPass signal and unusual local PCA geometry.

Windows with more distinctive local population structure also tended to have weakly elevated within-population LD (Figure 5C). The correlation between lostruct-score percentile and the median population-specific 5–50 kb LD percentile was positive but small in both NWA (Spearman’s rho = 0.083, circular-permutation p = 0.001) and NEP (rho = 0.127, p = 0.001). This supports increased local linkage over intermediate distances in at least some windows with high lostruct-score ranks, but elevated LD alone does not establish structural variation or locally suppressed recombination.

### Chromosome 6 as an illustrative region of distinctive local population structure

In the NEP lineage, BayPass revealed an approximately 0.8 Mb interval of elevated GEA signal at Bs6:3.145–3.950 Mb (Figure 3A). Local PCA across this interval revealed three discrete clusters of individuals (Figure 5D). The separation was clearest in the first portion of the interval and dissipated in flanking regions, providing a concrete example of the localized population-structure patterns summarized by the genome-wide lostruct analysis.

The frequency of the three local-PCA classes varied strongly among populations and along the NEP coast. However, the intermediate class did not show the excess heterozygosity expected under a simple two-arrangement inversion model. Within Bs6:3.145–3.350 Mb, mean PCAngsd F was 0.024 in the southern outer class (n = 17), -0.004 in the intermediate class (n = 36), and -0.029 in the northern outer class (n = 170). Exploratory within-population LD analyses likewise did not reveal a decisive elevation across the interval (data not shown). We therefore treat this region as an illustrative example of geographically structured local covariance rather than a confirmed inversion or heterokaryotype system.

### Spatial clustering of distinctive local-structure windows

Candidate windows were spatially clustered in several parts of the genome rather than occurring as independent, uniformly distributed observations. The full lostruct-score tracks and joint lostruct-PC1 + PC2 calls are shown together in the Supporting Information. This clustering motivated the chromosome-wise circular null used in the formal enrichment tests. Because those tests were performed on unmerged 100 kb windows, descriptive grouping of adjacent windows did not affect the genome-wide conclusions.

## DISCUSSION

Across two strongly differentiated North American lineages of *Botryllus schlosseri*, genomic variation was associated with broad thermal gradients across thousands of loci, consistent with repeated local adaptation to temperature. This pattern was nonetheless only partially shared between lineages. Most significant SNPs, genomic windows, and genes were unique to one lineage, although cross-lineage concordance and overlap exceeded null expectations at multiple genomic scales. GEA signal was only modestly associated with genomic regions displaying atypical patterns of local population structure, and these regions did not disproportionately contain the parallel component of the response. Together, these patterns suggest that local adaptation to temperature may proceed rapidly in this marine invader, while the particular variants and genomic regions involved remain strongly contingent on lineage history and genetic background.

### Repeated genomic responses across divergent thermal gradients

The NWA and NEP lineages provide unusually distinct genomic backgrounds across which to evaluate repeatability. Mean pairwise F_ST_ between populations from the two lineages was approximately 0.19, and substantial structure was observed within lineages. Nonetheless, nucleotide diversity remained high within individual populations, consistent with observations of extraordinary levels of genetic diversity in other tunicates (Leffler et al., 2012; Tsagkogeorga et al., 2012). Thus, the two coasts are not simply replicated environmental transects drawing on a shared, well-mixed pool of variation. Rather, selection appears to act on abundant variation that is strongly partitioned among populations and lineages. Against this background of pronounced population structure, the large number of temperature-associated loci in both NWA and NEP is consistent with extensive genomic differentiation along the thermal gradients on either coast. This pattern complements previous physiological work showing latitudinal differences in heat and cold tolerance on both coasts (Tobias et al., 2024, 2025). Although neither the phenotypes measured in wild-collected individuals nor the genomic associations presented here definitively establish rapid thermal adaptation on their own, together they support a model in which evolutionary adaptation contributes to invasion success in *B. schlosseri*. More broadly, the concordance of phenotypic and genomic differentiation across independent ranges supports the idea that environmental gradients encountered during range expansion can generate substantial evolutionary responses on contemporary timescales.

The degree of molecular repeatability depended on the scale at which the response was viewed. Exact SNP-level reuse was limited: only 161 SNPs were significant in both lineages, and same-sign effects were only modestly enriched. Even at this fine scale, however, the magnitude-weighted tests indicated that loci with larger temperature-associated effects tended to shift more concordantly across lineages than expected. Repeatability became clearer when nearby variants or different variants within the same gene were considered to represent a common response. Significant 5 kb windows overlapped more often than expected under both SNP-density-matched and chromosome-wise circular null models, and genes containing significant exonic variants also showed excess overlap. Window-level overlap was observed at 500 bp but not at 50 kb under the circular null, indicating that repeatability was concentrated at relatively fine genomic scales. This hierarchy is consistent with the view that parallel evolution is a continuum whose apparent strength depends on biological scale rather than a binary property of a response (Bolnick et al., 2018). A similar pattern has been reported across parallel temperature gradients in eelgrass, where SNP-level overlap was minimal despite repeatability at the gene level and evidence for shared polygenic responses across replicate bays (Schiebelhut et al., 2023). A recent study of island scrub-jays likewise found little parallelism at the SNP or gene level but substantial convergence at the pathway level, further suggesting that repeated adaptation can become more apparent at higher levels of biological organization (Cheek et al., 2026). In these examples, similar environmental pressures appear capable of producing a repeated genomic response without requiring extensive reuse of the identical nucleotide variants.

This predominance of lineage-specific associations in rapid adaptation is expected if evolutionary replicates begin with different pools of standing genetic variation. Theory and empirical studies increasingly show that the probability of molecular reuse declines as the genetic backgrounds of replicate populations diverge, because the same beneficial alleles are less likely to be available in each lineage and because genetically redundant routes can generate similar organismal responses (Bollback & Huelsenbeck, 2009; Bolnick et al., 2018; Conte et al., 2012). Across *Arabidopsis* and its relatives, for example, reuse of the same genomic basis of alpine adaptation declined with increasing divergence among lineages (Bohutínská et al., 2021), while experimental evolution has similarly shown declining parallel gene-expression responses with increasing genetic divergence under common selection (Thorhölludottir et al., 2025). The combination of high within-population diversity and strong differentiation in *B. schlosseri* fits this framework well: selection has abundant variation to draw on, but it is heavily partitioned among lineages. This makes the predominance of lineage-specific GEA unsurprising, while also making the excess cross-lineage overlap particularly notable. The evolutionary origins of those repeated signals remain unresolved with our low-coverage data, and could reflect retained ancestral variation, recurrent mutation, or different markers linked to the same causal locus.

This mixture of repeatability and contingency is particularly relevant to forecasting evolutionary responses to global environmental change. Genomic-offset approaches increasingly use present-day genotype-environment associations to project future maladaptation, but their predictive performance can depend strongly on demographic history, genetic architecture, and the extent to which contemporary adaptive relationships are transferable among populations INSERT DISCUSSION OF LIND et al 2025 here. The predominance of lineage-specific GEA in *B. schlosseri* highlights this limitation: similar environmental gradients can produce partially repeatable genomic responses without requiring the same variants or genomic regions to be involved. For widespread species facing rapid climate change, adaptive responses may therefore be more predictable at the level of environmental response than at the level of individual loci.

This mixture of repeatability and contingency is particularly relevant to forecasting evolutionary responses to global environmental change. Genomic-offset approaches increasingly use present-day genotype-environment associations to project future maladaptation, but their predictive performance depends on the evolutionary and environmental context in which those relationships were established (Fitzpatrick & Keller, 2015; Rellstab et al., 2021). In a simulation study, Lind & Lotterhos (2025) found that genomic-offset performance varied with demographic and adaptive context and declined when predictions extrapolated beyond the environments used for model training. Empirical tests likewise show variable transferability: genomic offsets have predicted biological responses in common-garden experiments of several tree species, but performance varies among environments, traits, and analytical approaches (Fitzpatrick et al., 2021; Lind et al., 2024; Verrico et al., 2026). In a recent marine common-garden experiment, genomic offsets predicted performance of eastern oysters only weakly and inconsistently across sites, further illustrating that predictive relationships aren’t guaranteed to transfer across contexts (Rumberger et al., 2026).

The predominance of lineage-specific GEA in *B. schlosseri* highlights a related limitation: similar environmental gradients can produce partially repeatable genomic responses without requiring the same variants or genomic regions to be involved. For widespread species facing rapid climate change, adaptive responses may therefore be more predictable at the level of environmental or phenotypic response than at the level of individual loci.

The functional annotations of the repeated candidates provide some additional biological context. Parallel exonic candidates included genes involved in mitochondrial respiration, redox homeostasis, cellular stress, environmental sensing, and metabolism. Among these, *SDHB*, which encodes the iron-sulfur subunit of mitochondrial complex II, is notable given longstanding hypotheses linking mitochondrial performance to thermal tolerance in ectotherms (Pörtner, 2001). Mitochondrial processes are strongly temperature-sensitive, and experimental work increasingly implicates maintenance of ATP synthesis, mitochondrial coupling, and redox balance in thermal performance (Chung & Schulte, 2020). Thus, repeated association at *SDHB*, together with candidates involved in oxidative stress such as *FTH1* and *GSTK1*, is consistent with selection on cellular energetic and redox physiology across thermal environments. Although far from definitive, these candidates provide plausible targets for future functional work.

Other repeated signals point more broadly to environmental sensing and organismal stress responses. *AHR*, for example, is a highly pleiotropic environmental-response regulator and has repeatedly contributed to adaptation to toxic pollutants in killifish (Reid et al., 2016). It also interacts with hypoxia signaling through the shared binding partner ARNT, linking xenobiotic sensing with oxygen-responsive transcription, although the nature of these interactions is context-dependent and not fully resolved (Chan et al., 1999; Vorrink & Domann, 2014). At a broader functional level, enrichment among gene-body candidates included immune and endocrine signaling, morphogenesis, and cell adhesion. Immune-associated enrichment is especially plausible along thermal gradients because temperature can alter both host immune performance and the abundance or virulence of pathogens; for example, Atlantic salmon show clinal MHC variation associated with temperature and bacterial diversity (Dionne et al., 2007). These results illustrate that thermal adaptation often involves contributions from multiple physiological systems. That pattern is consistent with a polygenic and diffuse architecture of thermal adaptation, with a modest repeated component embedded within a predominantly lineage-specific genomic context.

### Restricted dispersal and the genomic architecture of repeated adaptation

The distribution of these associations across the genome addresses a second question raised by *B. schlosseri*’s dispersal biology. Under migration-selection balance, gene flow can oppose local adaptation by continually introducing maladaptive alleles and, when recombination is appreciable, by breaking apart locally favored multilocus combinations. Theory therefore predicts that the selective advantage of clustering adaptive alleles should depend jointly on migration and recombination: as gene flow increases, selection may increasingly favor tighter linkage, larger-effect loci, or genomic rearrangements that reduce recombination among locally favored variants (Tigano & Friesen, 2016; Yeaman, 2013; Yeaman & Whitlock, 2011). Comparative empirical evidence is consistent with this expectation. Across six riverine fishes, Shi et al. (2023) found increasingly clustered distributions of adaptive outliers as inferred gene flow increased; the two most differentiated species showed relatively diffuse outlier distributions, whereas nearly all outliers in the least differentiated species occurred in a single large putative inversion.

*B. schlosseri* occupies a very different position along this migration-recombination continuum. Routine larval dispersal is extremely limited (Grosberg, 1987), mean pairwise F_ST_ even within focal lineages is approximately 0.1 (Figure 2), and within-population LD generally decays over roughly one kilobase. This rapid LD decay is also consistent with the unusually high recombination rates reported in other tunicates, such as *Ciona intestinalis* (Kano et al., 2006). This combination of weak homogenizing migration and rapid breakdown of linkage should reduce the selective advantage of concentrating locally adaptive variation into large, recombination-suppressed blocks. Under such a demographic and genomic regime, locally favored alleles may instead increase and persist more independently across the genome.

Our genome-wide analyses broadly support that expectation, although not universally. Temperature-associated variation tended to occur slightly more often in regions with distinctive local population structure, particularly in NEP, and these regions also showed slightly elevated within-population LD. However, the overall relationship was modest, and most GEA, including the parallel component, was not concentrated in these regions. We therefore interpret the absence of strong enrichment cautiously, as local covariance and LD provide only indirect evidence about underlying recombination or structural variation. Overall, extended regions of distinctive local population structure appear to contribute to some lineage-specific associations, but they do not seem to dominate the repeated genomic response.

This differs from several invasive systems in which large linked haplotypes or structural variants contain a disproportionate share of climate-associated or repeated adaptive variation. In common ragweed, large haploblocks representing about one tenth of the genome contained roughly one quarter of genomic windows showing parallel climate adaptation between native and invasive ranges (Battlay et al., 2023). In the globally invasive white clover, five large haploblocks demonstrated highly signficant, parallel climate-associated signals across independently colonized ranges; field experiments demonstrated strong fitness effects for three of them (Battlay et al., 2025). In the marine realm, rapid thermal adaptation in invasive European green crab (*Carcinus maenas*) has been associated with a single large, strongly linked genomic region, likely representing an inversion, whose allele frequencies track temperature and cold tolerance across both native and independently colonized ranges (Tepolt et al., 2022; Tepolt & Palumbi, 2020). Importantly, however, prominent structural variants need not imply that adaptation is dominated by them. In *Littorina saxatilis*, inversions contributed substantially to repeated ecotype divergence, but polygenic variation outside inversions made a contribution of similar magnitude (Koch et al., 2022). Likewise, although haploblocks were disproportionately involved in parallel climate adaptation in common ragweed, the majority of parallel genomic windows occurred outside these regions (Battlay et al., 2023, 2025). Thus, large-effect haploblocks and distributed adaptive variation can contribute simultaneously to repeated adaptation, with their relative importance varying among systems. The *B. schlosseri* results appear to fall toward the more distributed end of this spectrum: local population structure varied markedly across the genome, but repeated thermal associations were not disproportionately confined to the most distinctive regions. In a species with little routine exchange among populations, the selective advantage of concentrating adaptive variation into a small number of shared haploblocks may simply be weaker than in systems where migration would otherwise continually erode local allele-frequency differences.

The stronger relationship between GEA and unusual local structure in NEP warrants additional caution, because it need not indicate either stronger selection or a greater abundance of structural variants. The contrast with NWA is complicated by a more complex invasion history there, including evidence for both putatively native and introduced mitochondrial haplotypes (Yund et al., 2015), whereas the NEP lineage has undergone a recent, stepwise northward range expansion. Serial founder events during expansion can generate allele surfing, extended ancestry tracts, and large allele-frequency clines even in the absence of spatially varying selection (Excoffier & Ray, 2008; Hallatschek & Nelson, 2008). Here, latitude, temperature, and the likely axis of range expansion are tightly correlated. This is an especially difficult setting for genotype-environment association analysis: methods that model neutral population covariance substantially reduce demographic confounding, but range expansion or isolation by distance aligned with an environmental gradient can still generate false-positive associations (Lotterhos & Whitlock, 2015; Rellstab et al., 2015). Our chromosome-wise circular tests address a related problem by preserving broad genomic clustering while randomizing its alignment with local-structure tracks, but they cannot determine why a particular region has unusual local covariance and cannot distinguish selection from allele surfing, recombination-rate heterogeneity, linked ancestry, or interactions among these processes.

The prominent chromosome 6 interval illustrates this ambiguity. Across ∼0.8 Mb, NEP populations show both a dense GEA peak and a striking three-class local-PCA pattern that varies geographically. Although superficially consistent with a large inversion or haploblock, neither heterozygosity nor exploratory LD analyses supported a simple inversion model. We therefore interpret the region as a strong local-structure outlier whose origin remains unresolved and could reflect selection, range-expansion history, ancestry, recombination-rate variation, or structural variation. Higher-depth and long-read genomic data will be needed to distinguish among these alternatives.

The observation of parallel GEA across these strongly differentiated lineages strengthens the interpretation that at least some of the shared associations are adaptive. Demographic processes can generate environment-correlated allele-frequency clines within a lineage, but they are less likely to produce excess correspondence at the same loci, windows, and genes in independently structured lineages. Thus, the repeated component provides an important form of internal replication, even though temperature itself cannot be isolated from all correlated environmental variables.

Taken together, our results show that genomic responses across parallel thermal gradients can be broadly repeatable while remaining predominantly lineage-specific and broadly distributed across the genome. High standing diversity may provide abundant material for local adaptation, while strong subdivision and restricted dispersal shape which variants are available in each lineage and reduce the need for adaptive alleles to be concentrated in large linked blocks. More broadly, these results emphasize that under weak routine migration, rapid local adaptation need not rely on the concentrated genomic architectures favored under strong migration–selection conflict; instead, the routes available to adaptation may depend critically on how dispersal, recombination, and standing variation interact across populations.

## ACKNOWLEDGMENTS

This work was funded by a National Defense Science and Engineering Graduate Fellowship to Z. Tobias, the Woods Hole Oceanographic Institution’s Ocean Venture Fund and Grassle Fund awards to Z. Tobias, and a North Pacific Research Board grant to C. Tepolt (award #2212). Many people aided in the collection of *B. schlosseri* specimens for this project, including Cynthia McKenzie, Terri Wells, Haley Lambert, Claudio DiBacco, J. Ben Lowen, Shon Schooler, Rachel Flannery, Thom Therriault, Calley Wasser, Brett Howard, Stuart Crawford, Gary Freitag, Marnie Chapman, and Angela Bowers. We would especially like to acknowledge the Haida Nation and the Lax Kw’alaams Band for facilitating access to specimens from their respective waters.

## DATA ACCESSIBILITY

All sequence data are deposited in NCBI SRA under BioProject PRJNA1237870. Code for summarizing environmental data, bioinformatic processing, and genomic analysis will be made public after manuscript acceptance.

## AUTHOR CONTRIBUTIONS

ZT: conceptualization, funding acquisition, investigation, methodology, visualization, writing – original draft

CKT: conceptualization, funding acquisition, supervision, writing – review & editing

## SUPPORTING INFORMATION

### EXTENDED METHODS

#### Collections and DNA extraction

Colonies were collected from 24 North American sites between August 2019 and September 2022. Collections were made by hand from the undersides of floating docks at recreational marinas and commercial wharves. We attempted to space sampled colonies by at least 2 m because the very short dispersal distance of *B. schlosseri* larvae makes neighboring colonies more likely to be close kin. Approximately 1 cm^2^ colony fragments were preserved in 95% ethanol, held on ice during transport, and subsequently stored at -80 °C.

Before extraction, individual zooids were dissected from the surrounding tunic with insulin syringes (Becton Dickinson, cat. no. 328289). Gut tissue and brooding embryos were removed. Tissue from 8-10 zooids from the same or adjacent systems of a colony was extracted with a DNeasy Blood and Tissue Kit (Qiagen, cat. no. 69506) following the manufacturer’s tissue protocol, except that DNA was eluted in 32 uL elution buffer.

DNA quantity was measured with a Qubit 3.0 fluorometer and dsDNA Broad Range assay (Invitrogen, cat. nos. Q33216 and Q32850).

#### Library preparation and sequencing

Libraries were prepared from 540 specimens following the low-input tagmentation approaches of Baym et al. (2015) and Therkildsen and Palumbi (2017), with the following modifications. All incubations were performed on a Bio-Rad T100 thermal cycler. For each library, 1-10 ng genomic DNA in 1 uL 10 mM Tris (pH 8.5) was combined with 0.25 uL TDE1 tagmentation enzyme and 1.25 uL TD buffer (Illumina, cat. no. 20034197) and incubated for 5 min at 55 °C.

We added Illumina adapters and unique dual indexes in two PCR stages. In the first stage, we added 1.25 uL IDT for Illumina Unique Dual Index reagent (Illumina, cat. no. 20027213) and 3.75 uL KAPA HiFi HotStart Library Amplification Mix (Roche, cat. no. 7958960001). Cycling conditions were 72 °C for 3 min; 98 °C for 2 min 45 s; eight cycles of 98 °C for 15 s, 62 °C for 30 s, and 72 °C for 3 min; followed by 72 °C for 1 min. For the second stage, we added 0.5 uL each of 10 uM primer P1 (AATGATACGGCGACCACCGA) and P2 (CAAGCAGAAGACGGCATACGA), plus 8.5 uL KAPA HiFi mix. Cycling conditions were 95 °C for 5 min; four cycles of 98 °C for 20 s, 62 °C for 20 s, and 72 °C for 2 min; followed by 72 °C for 2 min.

Each reaction was brought to 30 uL with 10 mM Tris (pH 8.5), cleaned with 30 uL KAPA Pure Beads (1:1 bead-to-sample ratio), washed twice in 80% ethanol, and eluted in 32 uL 10 mM Tris. Library quantity was measured with a Qubit dsDNA High Sensitivity assay (Invitrogen, cat. no. Q32854). Equal DNA masses were combined into pools of less than 400 uL, precipitated with 0.1 volumes of 3 M sodium acetate (pH 5.2) and 2.5 volumes ice-cold ethanol, and incubated overnight at -80 °C. Pools were centrifuged at 14,000 rpm for 20 min at 2 °C, washed twice with 70% ethanol, air dried for 5-10 min, and resuspended in 32 uL 10 mM Tris.

Rather than bead-based size selection, concentrated pools were size selected on a Blue Pippin using a 1.5% agarose cassette in range mode, retaining fragments from 500 to 8,000 bp. Fragment distributions were checked on a Fragment Analyzer at the MIT BioMicroCenter. Pools were combined at equimolar concentrations and sequenced across six lanes of three NovaSeq 6000 SP flow cells (150-bp paired-end reads) at the University of Utah High-throughput Genomics Center.

#### Read processing and analytical sample set

Raw reads were inspected before and after trimming with FastQC v0.12.1 and summaries were collated with MultiQC v1.27.1. Trim Galore v0.6.10 was run in paired-end Nextera mode with Phred+33 encoding, a quality threshold of 20, adapter-stringency 2, and a minimum retained read length of 20 bp. Reads were mapped to the primary chromosome-level *B. schlosseri* assembly of De Thier et al. (2025) with Bowtie2 v2.5.4 in --very-sensitive-local mode. Mapping used forward-reverse orientation, a maximum insert size of 1,500 bp, and sample-specific read-group tags.

SAM files were converted and sorted with SAMtools. We retained reads with mapping quality at least 20, separated concordant, discordant, and singleton mappings, and carried only clipped concordant reads plus singletons forward to the final BAM. Overlapping segments of concordantly mapped read pairs were soft-clipped with bamUtil v1.0.15; reads rendered unmapped by clipping were removed. PCR duplicates were removed with Picard MarkDuplicates v2.26.3 (REMOVE_DUPLICATES=true), and reads were realigned around indels with GATK v3.8.1 RealignerTargetCreator and IndelRealigner.

One library (KWLM_3) had no assigned reads. QUIV_13 had extremely low mapping to the reference and mitochondrial sequence identified it as *Botrylloides violaceus*; it was excluded. An initial ANGSD coverage and missingness screen then identified six further samples site-level missingness greater than 80% (MSST_13, QUIV_26, RICH_28, SABS_23, SABS_2, and SABS_7). The retained all-sample dataset therefore contained 532 *B. schlosseri* individuals. Supplementary Figure S1 shows the distribution of per-individual mean coverage and proportion of the genome with coverage by population.

#### Genotype likelihoods, site filtering, and population assignments

Analyses used genotype likelihoods rather than hard genotype calls. ANGSD v0.938 was run in a Singularity v3.7 container retrieved from https://github.com/James-S-Santangelo/singularity-recipes/blob/master/angsd/Singularity.angsd_v0.938. Genotype likelihoods were computed using the SAMtools likelihood model (-GL 1) with BAQ adjustment (-baq 2) and retaining uniquely mapping, proper pairs after removal of bad reads. The genome-wide second pass was conducted independently for each of the 16 reference chromosomes. It required a base and mapping quality of at least 20, excluded triallelic sites, used an SNP likelihood threshold of 1e-6, required data from at least half of retained individuals, and imposed a total-depth maximum equal to 2.98 times the number of samples. The latter threshold was based on the first-pass depth distribution (three times the 90th percentile of mean sample depth) and was intended to limit repetitive or paralogous signal. The output Beagle genotype-likelihood and minor-allele-frequency files formed the basis for broad population-structure analyses.

We also evaluated a more stringent suite of SNPs, requiring representation in at least 90% of individuals as a missingness sensitivity analysis. The major global clusters recovered by PCA (see below) were unchanged (Supplementary Figure S2), indicating that the principal structure was not an artifact of differential site missingness. Minor-allele-frequency files were filtered to MAF ≥ 0.05, pre-thinned to a minimum 100-bp spacing, and analyzed with ngsLD v1.2.0 for pairs separated by ≤50 kb. prune_graph was then used to select an approximately independent subset, treating r2 ≥0.2 as linked. PCAngsd v1.03 was applied to the resulting genotype likelihoods. The same lineage-specific workflow was repeated within the major clusters to verify finer population assignments (Supplementary Figure S3).

Inspection of the NWA PCA identified four DART individuals (DART_2, DART_3, DART_4, and DART_19) that separated from the primary DART cluster. Because these may represent mislabeled individuals or recent migrants, we retained them for lineage-wide analyses but removed them from within-population analyses: population allele-frequency estimation, site-frequency spectra, F_ST_, nucleotide diversity, within-population LD, and BayPass input construction. Thus, the NEP and NWA population-level datasets contain 10 and 12 populations, respectively, whereas global analyses retain the full lineage sample set.

#### Population differentiation, within-population LD, and nucleotide diversity

For pairwise F_ST_, we estimated folded two-dimensional site-frequency spectra with winSFS v0.7.0 from population-specific genotype-likelihood data. The folded spectra were supplied to realSFS fst index and fst stats to estimate pairwise values. Folding avoids reliance on ancestral-state polarization and is appropriate for the reference-based data used here.

The within-population LD analysis was deliberately separated from the lineage-wide pruning calculations, because pooling highly structured populations can create long-distance statistical LD unrelated to physical linkage. For each focal NWA and NEP population, we generated genotype likelihoods after filtering to MAF ≥0.05 and pre-thinning sites to a minimum 100-bp separation. ngsLD was run separately for every population and chromosome with posterior genotype probabilities and a 50-kb maximum pair distance. We summarized the genotype-likelihood-based r2_ExpG statistic in 100-bp distance bins. The reported decay position is the first of ten consecutive bins whose mean r2_ExpG was ≤0.2. Requiring a full 1-kb run of bins makes this an intentionally conservative estimate in the noisy low-coverage data; it is not a fitted recombination-rate estimate.

For nucleotide diversity, each population was evaluated at the same 204,842,689 analyzed sites. Folded one-dimensional SFSs were estimated with winSFS and passed, without the winSFS metadata header, to realSFS saf2theta with -fold 1. thetaStat provided one pairwise-theta summary per chromosome. Genome-wide pi was the sum of pairwise theta (tP) across the 16 chromosomes divided by the common number of analyzed sites. This use of realSFS calculates theta from an SFS estimated by winSFS; it does not use realSFS to estimate the SFS. Per-population values are reported in Table S2.

#### Environmental temperature data extraction and provenance

Environmental data were assembled before GEA analysis from *in situ* water-temperature records rather than gridded products. This is more appropriate for our *B. schlosseri* populations, which were collected and live their entire lives in the shallow subtital on coastal infrastructure. Table S1 gives the sampling coordinates, station or report used, station coordinates, approximate distance from the sampling site, record dates, source URL, and final mean annual temperature (MAT). Raw downloads and the processing notebook are archived in the project directory "Station Temp Data/for GEA".

For the standard workflow, raw station tables were harmonized to timestamp and temperature fields. Temperatures outside -5 to 40 °C were removed. Observations were averaged within day; daily values were then averaged by calendar date across years; and MAT was the mean of this day-of-year climatology. This gives each day of the annual cycle equal weight rather than giving extra weight to days sampled more frequently. Where data permitted, the target interval was 1 January 2015 to 19 October 2023. Actual observation intervals differ among stations because some records began later or ended earlier; these intervals are reported explicitly in Table S1 rather than imputed.

Most populations used local or regional NOAA, NOAA National Estuarine Research Reserve System, Smart Atlantic, USGS, SCCOOS, or comparable instrumental records. Four sites required more explicit decisions:

1. KWLM (Lax Kw’alaams) used the Ocean Networks Canada Prince Rupert Atlin Terminal Station. The 6-m record was restricted to 1 July 2017-19 October 2023 to exclude an isolated early artifact. Daily means over this interval gave MAT = 9.02 °C.
2. MSST (Masset) had no suitably local sheltered instrumental record. We used the annual mean at Fisheries and Oceans Canada’s Langara Light Station, which is more exposed, and added a 0.51 °C Masset-minus-Langara offset estimated from older paired observations. This yielded MAT = 9.90 °C.
3. NCSL (Newcastle Island) used the nearby, sheltered Departure Bay Light Station directly. Its record had few gaps and gave MAT = 12.06 °C.
4. REED (Port Moody) was represented by a regional Burrard Inlet physical-parameters report. The available Ocean Networks Canada sensors were deeper (approximately 22-39 m), farther seaward, and did not match the report’s interval, so they were inspected but not used. MAT was the unweighted mean of the report’s four seasonal values: (8.0 + 9.6 + 17.7 + 11.2)/4 = 11.625 °C, reported as 11.63 °C.

The complete provenance table (Table S1) is intended to make the proxy decisions transparent and reproducible. We initially calculated MAT, mean winter temperature, mean summer temperature, mean number of days ≥12 °C, and latitude. These variables were strongly collinear across the sampled populations, so mean annual temperature was retained as the single BayPass covariate. It remains a biologically motivated summary of the thermal environment, but it cannot be separated completely from latitude and correlated coastal environmental gradients in this design.

#### BayPass input construction and GEA models

ANGSD was run separately by population to estimate allele frequencies and obtain site-level read depth. These population-level runs used the SAMtools likelihood model, BAQ adjustment, removal of bad and non-unique reads, proper-pair filtering, base and mapping quality ≥30, and a within-population minimum-individual threshold of approximately 25% of samples. The maximum total-depth threshold was scaled to approximately 2.9 times the number of individuals in that population.

BayPass inputs were constructed as integer pseudo-counts rather than raw read-base counts. For each population and site, the ANGSD knownEM allele-frequency estimate was multiplied by the number of called chromosomes (2N) and rounded to obtain one allele pseudo-count; the complementary pseudo-count was 2N minus that value. Pseudo-count pairs were then reconciled to a common allele orientation within each lineage. At each site, both pseudo-counts for a population were set to zero when its total read depth was <20. A site was retained when it was represented in at least 10 of 12 NWA populations or at least 9 of 10 NEP populations, then filtered to lineage-wide MAF ≥0.05. These criteria yielded 8,770,990 NWA and 8,858,634 NEP sites. The masking step ensures that a nominally analyzed population does not contribute uninformative low-depth counts to BayPass.

We estimated the BayPass omega covariance matrix from an LD-pruned subset of the final pseudo-count matrices. Candidate sites were pre-thinned to 100-bp spacing, ngsLD was run within chromosomes up to 50 kb, and prune_graph removed sites with r2 ≥0.2. BayPass v2.1 neutral-model runs were repeated across five seeds for each lineage. The resulting omega matrices were highly similar; a representative omega matrix was used for the corresponding GEA runs. The complete pseudo-count and site files were distributed into 200 interleaved chunks for computational tractability. Each chunk was analyzed with BayPass -covmcmc and -scalecov using lineage-specific omega and MAT, and the complete procedure was repeated for five seeds.

The BayPass summary-betai output supplies both eBPmc, the empirical Bayesian association score, and M_Beta, the posterior-mean regression coefficient for MAT. For every SNP, we took the median eBPmc across the five GEA replicates. SNPs with median eBPmc ≥3 were classified as GEA-positive. This is the reported threshold corresponding to an empirical Bayesian p-value <0.001. M_Beta values were retained for the SNP-level parallelism analysis; no separate absolute-M_Beta cutoff was applied.

#### Parallelism tests

All cross-lineage tests were restricted to locations represented in both BayPass datasets. For shared loci, allele orientation was harmonized across lineages before comparing M_Beta, so effect signs always referred to the same allele. The analyses intentionally treat parallelism as a scale-dependent property rather than requiring the same nucleotide to be an outlier in both lineages.

#### SNP-level direction and magnitude

For each ascertainment direction, we retained GEA-positive SNPs in the focal lineage, assigned them to 50-kb bins, and sampled one SNP per occupied bin in each replicate. This broad thinning prevents a localized cluster of non-independent signals from dominating a resample; it is deliberately more conservative than the approximately 1-kb population-level LD scale. For each draw, we calculated (i) the fraction of sampled SNPs whose M_Beta had the same sign in both lineages and (ii) the mean cross-lineage product of M_Beta. The null was generated by randomizing the signs of one lineage’s M_Beta values. This preserves the SNP set, spatial thinning, and absolute effect-size distribution. Positive products arise from same-sign effects and negative products from opposite-sign effects; near-zero effects make little contribution. The product statistic therefore evaluates magnitude-weighted directional concordance without creating an arbitrary effect-size threshold.

#### Window-level overlap

We tiled each chromosome into non-overlapping 500-bp, 5-kb, and 50-kb windows. The focal scale was 5 kb, with the other scales treated as sensitivity analyses. At each scale, we used two summaries: (i) a maximum-eBPmc definition, under which a window was positive if it contained at least one eBPmc ≥3 SNP; and (ii) a mean-eBPmc definition, under which windows in the lineage-specific upper 5% of mean eBPmc were positive. A parallel window met the relevant definition in both lineages. The analyses were restricted to windows with at least one BayPass SNP in each lineage.

The matched window null controls heterogeneous opportunity to detect an association. Within each chromosome, windows were binned by the deciles of their SNP counts in NWA and NEP, and lineage labels were permuted within these joint bins before recalculating overlap. The chromosome-wise circular null addresses a separate issue: broad spatial clustering. The positive-window track of one lineage was shifted by a random offset within each chromosome, wrapping at chromosome ends, before overlap was calculated. It therefore preserves the number and broad spatial structure of outlier windows on each chromosome while randomizing their alignment between lineages. Each null used 1,000 permutations; one-sided empirical p-values were calculated with a +1 correction.

#### Gene-level overlap

We reduced overlapping exons across transcripts within each annotated gene before assigning BayPass SNPs, thereby avoiding duplicate exon bases and duplicate SNP-gene assignments. The conservative gene analysis considered only exonic SNPs. A gene was positive if it contained at least one eBPmc ≥3 exonic SNP; the sensitivity definition selected the upper 5% of genes ranked by mean exonic eBPmc. The background comprised genes with at least one exonic BayPass SNP in both lineages. For the significant-SNP test, null gene sets were sampled independently within deciles of reduced exon length. For the mean-eBPmc test, they were sampled within joint deciles of the NWA and NEP exonic SNP counts. These matched nulls account for the greater opportunity for long or densely assayed genes to contain an outlier. Each test used 1,000 permutations and the same one-sided empirical p-value convention.

#### Functional annotation and enrichment

Predicted proteins were annotated with eggNOG-mapper v2.1.13 against eggNOG v5.0.2 using DIAMOND v2.0.15. Orthology-based annotations with e-values <1e-10 were retained. Table S3 lists the high-confidence functional annotations for the 113 conservative, parallel exonic candidate genes that received an eggNOG orthology assignment.

The functional-enrichment analysis intentionally used a broader gene-body candidate set: genes containing at least one significant BayPass SNP anywhere in the annotated gene body in both lineages. This permits putatively regulatory associations to contribute, whereas the conservative annotation table is limited to exonic candidates. We tested GO-term over-representation with clusterProfiler against the background of genes containing at least one analyzed SNP in both lineages. Significant terms across the three GO namespaces were clustered using Jaccard overlap among their member-gene sets. Figure 4 displays the readable cluster-level summary; Table S4 reports the individual significant GO terms, their nominal and adjusted p-values, and the number of contributing genes.

#### Extended regions of distinctive local population structure

We used local PCA and lostruct v0.0.0.9000 to describe genomic heterogeneity in local population structure. For each lineage, PCAngsd estimated a covariance matrix from genotype likelihoods in non-overlapping 100-kb windows. A window was run when it contained at least 10 input sites; for the lostruct analysis, we retained only windows with at least 250 sites retained by PCAngsd after MAF filtering. For each retained covariance matrix, we calculated the eigendecomposition and supplied the total sum of squares, first two eigenvalues, and first two eigenvectors to lostruct. lostruct::pc_dist compared local-PC representations using L1 normalization, and classical multidimensional scaling was applied to the inter-window distance matrix. We retained MDS1-MDS5.

For a continuous score, we calculated the median coordinate across all windows for each MDS axis within a lineage and then calculated each window’s Euclidean distance from this five-dimensional median center. The original MDS scale was retained, so axes contributing more strongly to between-window distance also contribute more strongly to the score. Scores were converted to within-lineage percentile ranks. A high score thus identifies a window with local PCA geometry unlike the lineage-wide background.

For the categorical analysis, we also calculated the proportion of local covariance explained jointly by PC1 and PC2. A window was designated as having distinctive, dominant-axis local population structure only if it fell in the top 10% for both the Euclidean lostruct score and PC1 + PC2 variance explained within its lineage. The common top-10% threshold makes the two components symmetric. Calls were made at the individual 100-kb-window level. Supplementary Figure S4 shows the continuous tracks and the individual joint calls, illustrating how window-level observations are spatially clustered.

#### Association of GEA with local population structure

We summarized GEA on the same 100-kb scale as local PCA by partitioning each local-PCA window into its constituent non-overlapping 5-kb windows. For each lineage, the response was the number of analyzable 5-kb windows containing at least one eBPmc ≥3 SNP out of the number of analyzable 5-kb windows. A parallel 5-kb window had at least one tested SNP and at least one significant SNP in both lineages; the significant SNPs themselves need not be identical. This definition matches the focal 5-kb window-parallelism analysis in the main text.

The continuous analysis used Spearman’s rho between the lineage-specific lostruct-score percentile and the proportion of GEA-positive 5-kb windows. For parallel GEA, the predictor was the larger of the NWA and NEP score percentiles, meaning unusual local population structure in either lineage. The categorical analysis used grouped-binomial logistic regression, with GEA-positive and GEA-negative 5-kb counts as the response and joint lostruct-PC1+PC2 candidate status as the predictor. The parallel categorical analysis used candidate status in either lineage. These models provide descriptive observed effect sizes; inference did not use their nominal model-based p-values.

For both continuous and categorical tests, we retained the lostruct track and rotated the GEA-positive and analyzable-bin counts together by circular permutation, using independent random offsets within each chromosome. This permutation approach preserves the amount of GEA signal, the denominator, and broad spatial autocorrelation within every chromosome, while disrupting their alignment with the fixed local-structure predictor. For each of 1,000 permutations, we recalculated rho or the log odds ratio. One-sided empirical p-values use [1 + number of permuted statistics ≥ observed statistic]/1,001. This is also why local-structure candidate windows were not merged for the formal tests.

The continuous association is not independent evidence for reduced recombination: selection or demography can affect both allele frequencies, and therefore BayPass signal, and local covariance. The categorical dominant-axis definition and the independent LD comparison below provide complementary, but still non-diagnostic, context.

#### Within-population LD and local population structure

To test whether windows with high lostruct-score ranks also had elevated linkage extending beyond ordinary short-range LD, we used population-specific ngsLD output rather than pooled-lineage LD. The full pairwise outputs were collapsed to one record per SNP containing the number of comparisons, mean r2_ExpG, and fraction of comparisons with r2_ExpG ≥0.2 in two distance classes: 0-5 kb and 5-50 kb. For every population and 100-kb local-PCA window, we calculated pair-count-weighted mean r2_ExpG. Because the input is SNP-centered, a pair contributes to the local LD environments of both endpoint SNPs; pairs spanning a window boundary can therefore contribute to both adjacent windows.

The primary response was the 5-50-kb class, which represents persistence beyond the approximately 1-kb background LD scale. To avoid conflating among-population differences in absolute LD with local enrichment, window summaries were converted to percentile ranks separately within each population and chromosome. We took the median percentile across populations within each lineage and retained windows represented by at least half of that lineage’s populations. We then used the same chromosome-wise circular-permutation framework as above, rotating the LD track within chromosomes while keeping lostruct scores fixed.

#### Illustrative NEP chromosome 6 region

The chromosome 6 analysis is included as an example of a localized region with distinctive population structure. We examined genotype likelihoods across Bs6:3,145,000-3,950,000 and adjacent intervals with PCAngsd. The three local-PCA classes were assigned descriptively from their positions in the local PC1-PC2 space. The clearest separation occurred in Bs6:3,145,000-3,350,000, so PCAngsd individual inbreeding coefficients were estimated jointly across individuals for that segment. We then summarized F by the two outer and intermediate local-PCA classes. Under a simple two-arrangement inversion model, the intermediate class would be expected to show substantially lower F (greater heterozygosity) than the outer classes. As reported in the main text, that pattern was not apparent. This result, together with the lack of a decisive within-population LD signature in exploratory analyses (data not shown), is why we describe the region as a geographically structured example of local covariance rather than a confirmed inversion or heterokaryotype system.

## SUPPLEMENTARY FIGURES

**Figure S1.**
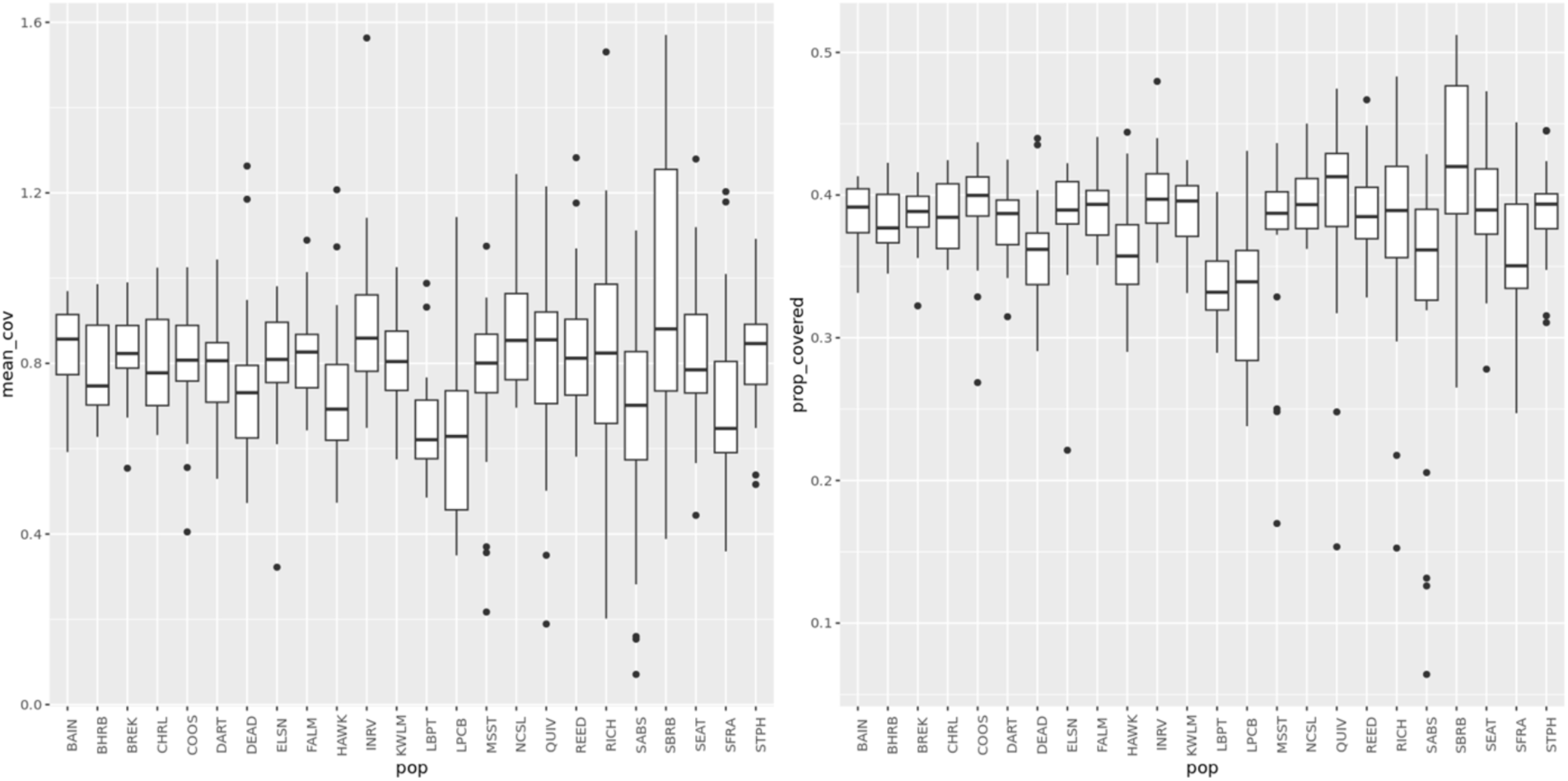
Distributions of mean coverage (left) and proportion of sites with data (right) across populations. Individuals with less than 20% of sites covered were excluded.

**Figure S2.**
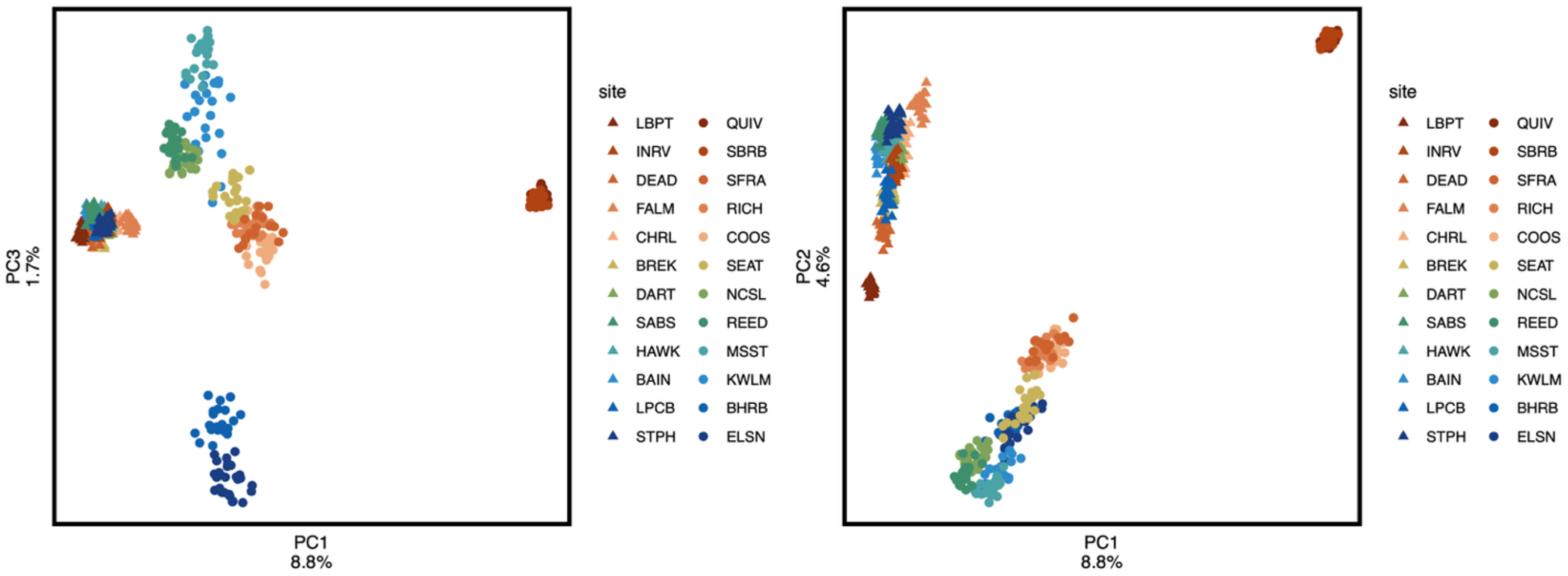
PCAs on subset of sites with data for at least 90% of individuals.

**Figure S3.**
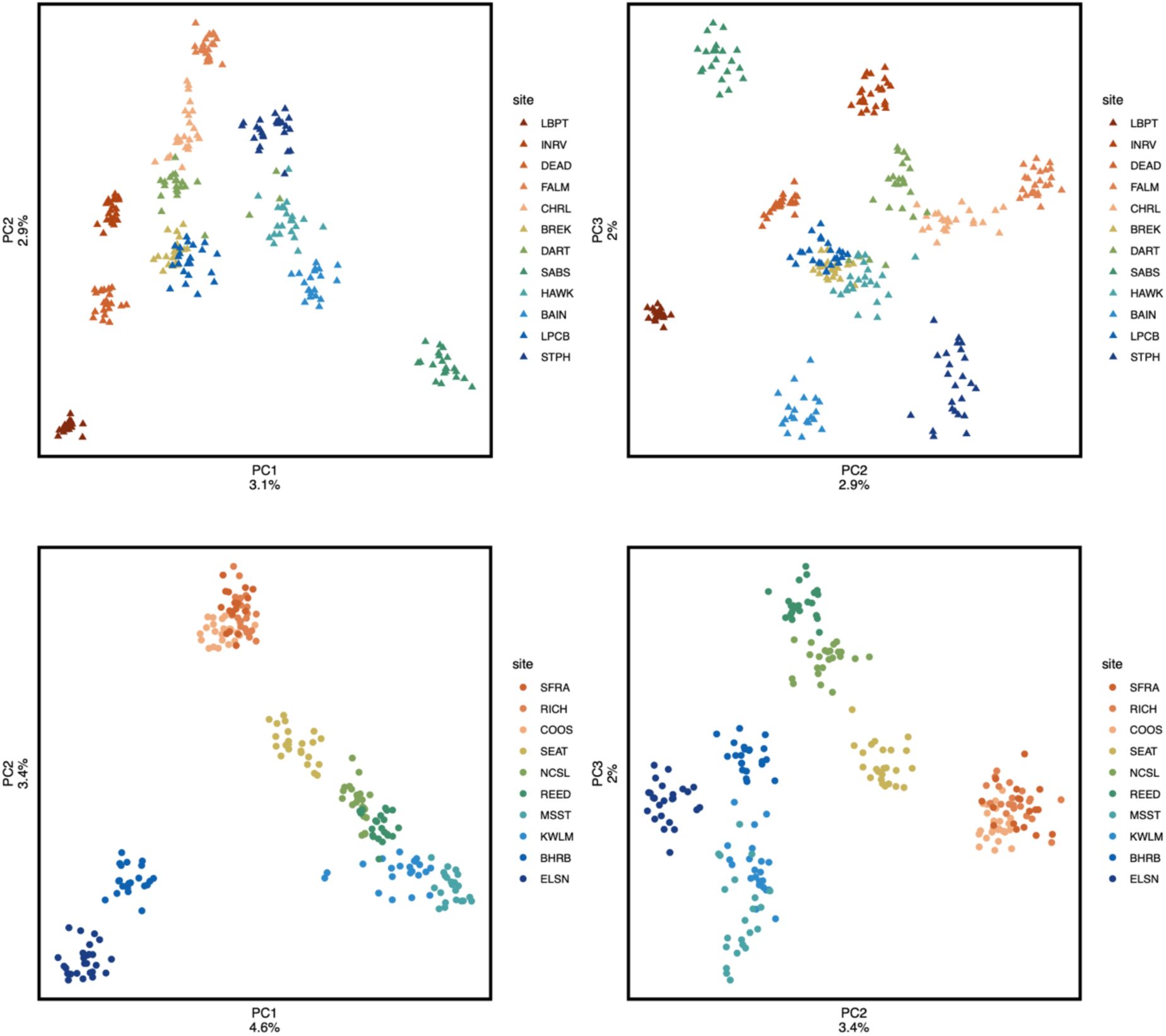
PCAs from genotype likelihoods estimated within each lineage demonstrate hierarchical population structure.

**Figure S4.**
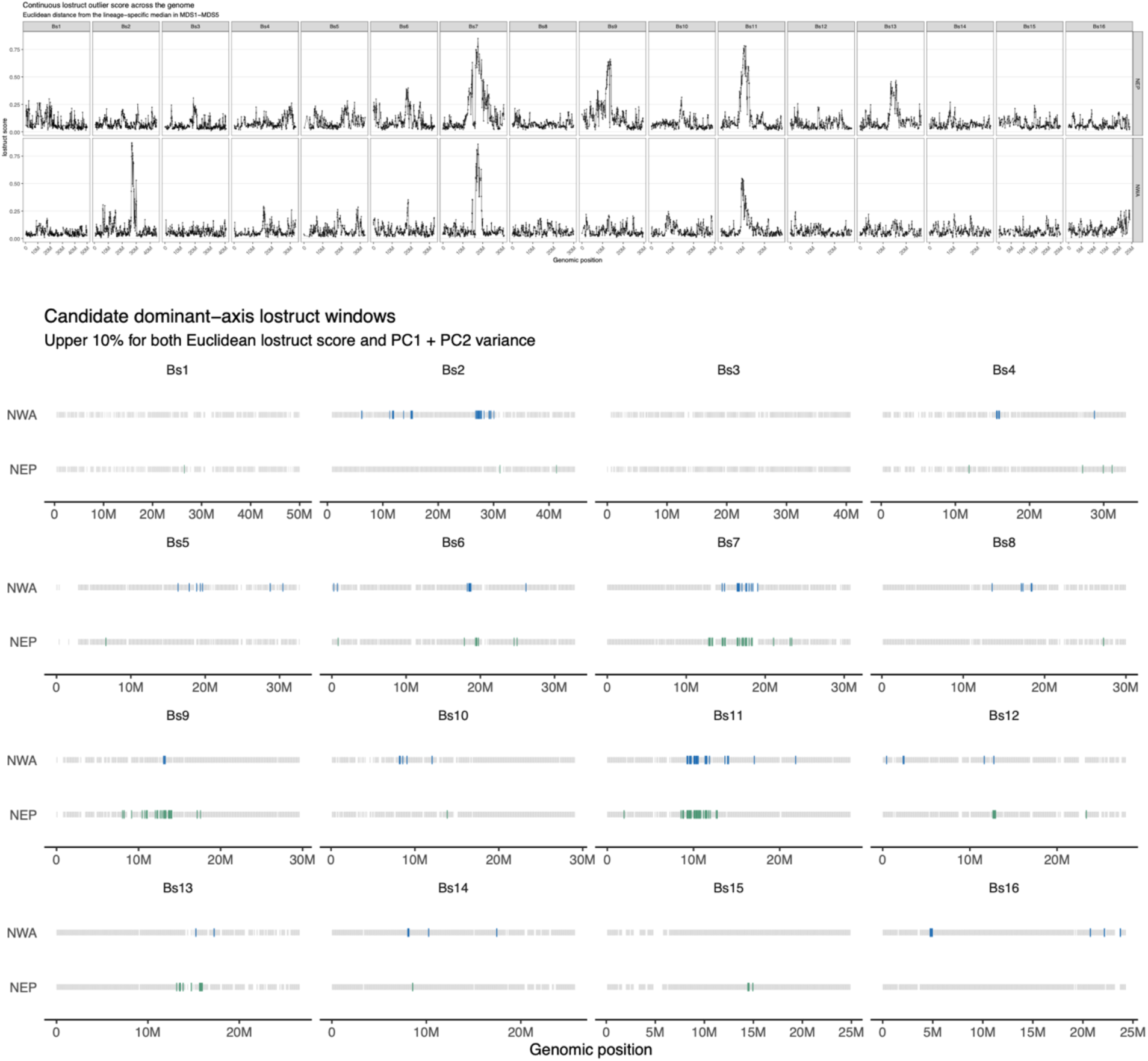
Distribution of Euclidean-distance lostruct score across genome for NEP and NWA (top). Regions in the top 10% of lostruct scores and top 10% of combined PC1+2 percent variation explained.

**Figure S5.**
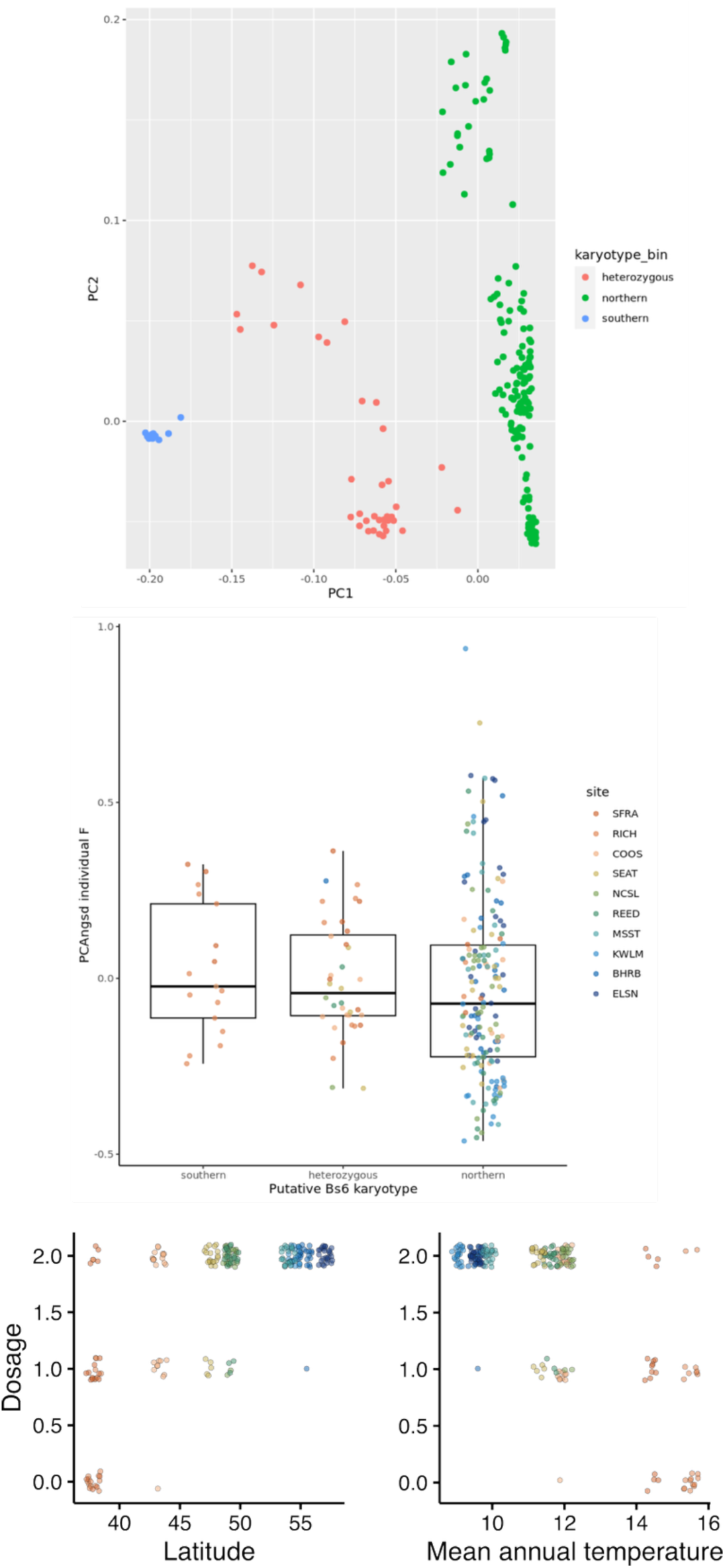
(top) PCA of NEP individuals the region on Bs6 from Figure 5. Points are colored by their pseudo-karyotype bin. (middle) Box plots of F stat distributions from PCAngsd for each of the three pseudo-karyotype bins. (bottom) Association with latitude (left) and mean annual temperature (right) based on number of “alleles” of this putative haploblock.

